# Mitochondrial damage triggers concerted degradation of negative regulators of neuronal autophagy

**DOI:** 10.1101/2024.06.12.598718

**Authors:** Bishal Basak, Erika L.F. Holzbaur

**Author notes:** **Correspondence** (E.L.F.H.).

## Abstract

Mutations in genes that regulate mitophagy, a key mitochondrial quality control pathway, are causative for neurological disorders including Parkinson’s. Here, we identify a novel stress response pathway activated by mitochondrial damage that regulates mitophagy in neurons. We find that increasing levels of mitochondrial stress triggers a graded, concerted response that induces proteasomal degradation of negative regulators of autophagy. These include Myotubularin-related phosphatase 5 (MTMR5), MTMR2 and Rubicon. This ‘**Mito**phagic **S**tress **R**esponse’ (MitoSR) pathway is neuron-specific and acts in parallel to the classical Pink1/Parkin-mediated mitophagy pathway. While MTMR5/MTMR2 inhibits autophagosome biogenesis, we find that Rubicon inhibits lysosomal function and thus blocks autophagosome maturation. Targeted depletion of these negative regulators is sufficient to enhance mitophagy, promoting autophagosome biogenesis and facilitating the fusion of mitophagosomes with lysosomes. Our work suggests that therapeutic activation of the MitoSR pathway to induce degradation of negative regulators of autophagy may enhance mitochondrial quality control in stressed neurons.

## INTRODUCTION

Mitochondria are the primary organelles responsible for fueling the cell’s metabolic needs by producing adenosine triphosphate (ATP)^1,2^. Neurons, by virtue of their polarized nature and high metabolic activity, are particularly dependent on mitochondria to supply energy to regulate neurotransmission, calcium homeostasis, and neural plasticity^3–5^. This elevated metabolic activity makes neuronal mitochondria more susceptible to damage; further, as neurons are post-mitotic, this damage can continue to accumulate over an organism’s life span. The build-up of toxic dysfunctional mitochondrial fragments is detrimental to neuronal health, as they can generate reactive oxygen species (ROS), pro-inflammatory signals and trigger apoptosis leading to neurodegeneration^6^.

Neurons employ different quality control mechanisms to prevent the accumulation of damaged mitochondria and hence preserve mitochondrial physiology^7,8^. One of the critical processes that regulates organelle turnover and homeostasis in eukaryotic cells is autophagy. Under basal conditions, neuronal autophagy initiates primarily at synaptic sites and at the axon terminal, where a growing phagophore engulfs nearby cytoplasmic components^9,10^. The autophagosome matures as it traffics retrogradely to the soma where its fuses with the acidic lysosome to mediate degradation of its cargo^9,11^.

Under basal conditions, mitochondria constitute ∼10-20% of the cargos degraded by non-selective autophagy in the brain^12^. However, when mitochondria undergo depolarization or oxidative stress, their targeted removal is mediated via receptor mediated selective autophagy called mitophagy^13–16^. In neurons, Pink1/Parkin mediated mitophagy remains the most well-characterized pathway for mitochondrial quality control by autophagy. In this pathway, the accumulation of PINK1 kinase on the outer mitochondrial membrane^17,18^ leads to the recruitment and activation of the E3 ligase Parkin^18–22^. Activated Parkin then ubiquitinates a plethora of mitochondrial proteins^23–29^. These ubiquitinated proteins serve as a binding platform for recruitment of autophagy receptors^30–38^ to promote the formation of an engulfing autophagosome that sequesters the damaged mitochondria away from the cytosol. Mutations in genes involved in this pathway lead to Parkinson’s disease (PD)^39,40^, Amyotrophic lateral sclerosis (ALS)^41,42^ and frontotemporal dementia (FTD)^43,44^. These observations suggest that mitophagy is a critically important pathway to maintain neuronal health and survivability; however, there is a limited understanding of how this process may be regulated in neurons.

Accumulating data suggest that the regulatory pathways that control autophagy in neurons are unique, as they are relatively resistant to autophagy inducers such as mTOR inhibition or amino acid starvation^45,46^. This resistance is partly mediated by the expression of myotubularin-related phosphatase 5 (MTMR5), a protein that negatively regulates autophagy in neurons^47^. MTMR5 belongs to the myotubularin family of proteins, which function as phosphatidylinositol 3-phosphate (PI3P) phosphatases. Proteins in this family such as MTMR6, MTMR8, MTMR9, MTMR14 have been reported to repress autophagy in different cell types^48–50^. MTMR5, a catalytically inactive member of the family, is particularly enriched in neurons^47^ and has been shown to modulate the localization and enzymatic activity of the active MTMR2 phosphatase^51^. The MTMR5-MTMR2 complex collectively represses early steps of autophagosome biogenesis by dephosphorylating PI3P produced by the action of VPS34^47^. Consistent with this model, knockdown of MTMR5 or MTMR2 in iPSC-derived neurons results in an increase in autophagosome number during mTOR inhibition^47^.

Another negative regulator known to down-regulate autophagy in non-neuronal cells is Rubicon^52,53^. Rubicon was first identified as a subunit of the class III phosphatidylinositol 3-kinase complex II (PI3KC3-C2). PI3KC3-C2 includes the subunits UVRAG, VPS34, VPS15 and BECN1, and plays critical roles in phagophore expansion and endosome maturation^54–56^. Rubicon associates with a subpopulation of PI3KC3-C2 to block recruitment of the complex to the growing membrane^57^. Consistent with this, overexpression of Rubicon in non-neuronal cells reduces PI3P formation^57^ and autophagosome number^52,53,57^. Rubicon also binds to Rab7-containing late endosomes and lysosomes and has been implicated in regulating membrane trafficking and reducing autophagic flux^52,53,58,59^. A recent study in HeLa cells revealed that phosphorylation of Rab7 upon mitochondrial damage induces the exchange of Rubicon for another Rab7-binding protein, Pacer, which is a positive regulator of autophagy. This exchange allows for a modest increase in mitochondrial turnover by autophagy following depolarization, supporting a role for Rubicon in repressing organellar homeostasis ^60^.

In the nervous system, recent findings indicate that systemic deletion of Rubicon in mice reduces the accumulation of phosphorylated α-synuclein upon injection of its native unphosphorylated form in the brain striatum^61^, a primary pathological hallmark of PD. Rubicon levels are also reported to be higher in the lumbar spinal cord region of ALS patients^62^. These studies thereby necessitate a detailed molecular understanding of the role of Rubicon in regulating neuronal homeostasis by autophagy.

Here, we demonstrate that MTMR5, MTMR2 and Rubicon are degraded as a graded, concerted response to increasing levels of mitochondrial damage in neurons. These results define a mitophagic stress response pathway, or MitoSR, that operates via the ubiquitin-proteasomal pathway and acts in parallel to the canonical Pink1/Parkin response. Activation of MitoSR occurs in response to multiple inducers of mitochondrial stress in neurons, but the pathway is not triggered upon lysosomal stress. Induction of MitoSR removes negative regulators of autophagosome biogenesis, MTMR5/MTMR2, and also targets Rubicon which we identify to block lysosomal function thereby inhibiting autophagosome maturation in neurons. Finally, we show that under mild mitochondrial stress, targeted depletion of the negative regulators of autophagy significantly increases mitochondrial turnover in neurons. Thus, we propose that therapeutic interventions directly targeting negative regulators of autophagy may promote clearance of damaged mitochondria in neurodegenerative diseases such as PD and ALS where mitophagy is compromised.

## RESULTS

### Mitochondrial stress initiates proteasomal degradation of negative regulators of autophagy

The process of autophagy is critical for neuronal survivability, yet neurons are largely resistant to major changes in basal autophagy flux. Recent progress has identified several negative regulators of autophagy, including MTMR5, MTMR2^47^ and Rubicon^53,55,57,60^ which can reduce autophagic flux in different cellular systems. This led us to ask under periods of cellular stress when there is a need to upregulate autophagy, can neurons bypass these autophagy suppressors. To test this idea, we treated mouse embryonic cortical neurons with increasing doses of Antimycin A (Ant A), a well-characterized mitochondrial depolarizing agent that inhibits complex III of the electron transport chain, generating localized ROS ^63,35,32,64^. Levels of MTMR5, MTMR2 and Rubicon remain unaltered under vehicle (ethanol-EtOH) treated condition or low doses (3 nM) of Ant A **(Figure 1A-D)**. However, with increasing Ant A concentrations (15 nM, 30 nM), we observed dramatic decreases in expression levels of MTMR5, MTMR2 and Rubicon. For MTMR2, we could also detect a parallel increase in the levels of a degraded product that migrates just beneath the intact protein on western blots. For Rubicon, we observed a doublet on western blots that may represent two isoforms **(Supplemental S1)**. However, the expression of both of these bands decreased in parallel in response to mitochondrial damage; hence we focused on the more prominent band in our analysis. Together, these observations indicate that there is a concerted decrease in the expression of three negative regulators of autophagy, MTMR5, MTMR2 and Rubicon, in response to increasing concentrations of Ant A. These findings suggest that mitochondrial ROS production triggers the dose-dependent degradation of these proteins.

**Figure 1:**
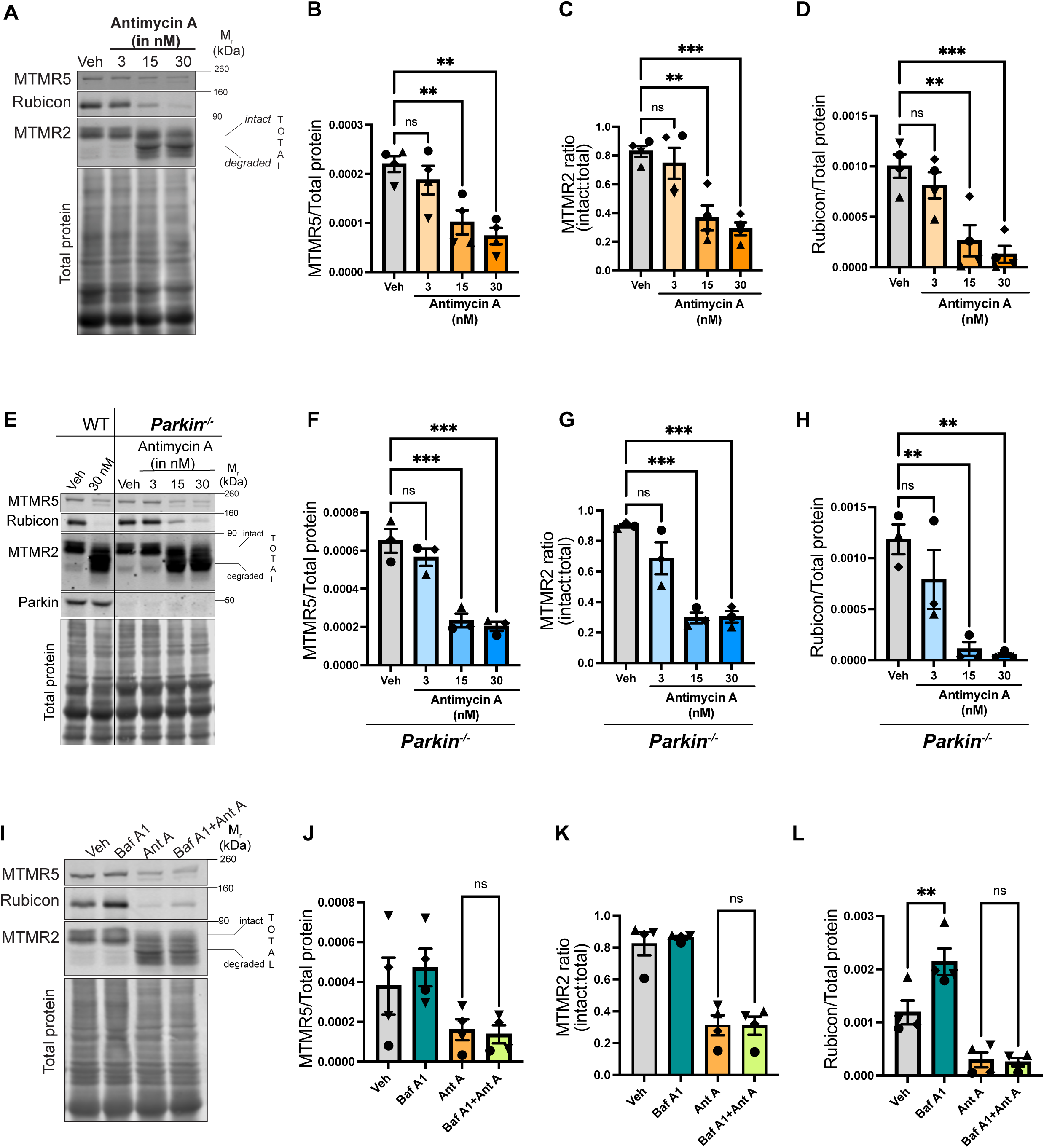
Mitochondrial stress initiates the concerted degradation of MTMR5, MTMR2 and Rubicon, independent of Parkin activity or autophagic flux. (A-D) MTMR5, MTMR2, and Rubicon are degraded in response to increasing concentrations of Ant A. (A) Representative western blots from lysates of wild type (WT) murine embryonic cortical neurons treated with vehicle (EtOH), or with increasing concentrations of Ant A (3 nM, 15 nM or 30 nM) for 2 hrs. (B) MTMR5 band intensity normalized to total protein from WT neurons treated with EtOH or increasing concentrations of Ant A (N=4 experiments, One way ANOVA with Dunnett’s multiple comparison test). (C) Ratio of intact MTMR2 band intensity normalized to total (intact + degraded) band intensities from WT neurons treated with EtOH or increasing concentrations of Ant A (N=4 experiments, One way ANOVA with Dunnett’s multiple comparison test). (D) Rubicon band intensity normalized to total protein from WT neurons treated with EtOH or increasing concentrations of Ant A (N=4 experiments, One way ANOVA with Dunnett’s multiple comparison test). (E-H) Concerted degradation of MTMR5, MTMR2, and Rubicon is also observed in cortical neurons from *Parkin^-/-^* mouse embryos. (E) Representative western blots from lysates of *Parkin^-/-^* murine embryonic cortical neurons treated with vehicle (EtOH), or with increasing concentrations of Ant A (3 nM, 15 nM or 30 nM) Ant A for 2 hrs. (F) MTMR5 band intensity normalized to total protein from *Parkin^-/-^* neurons treated with EtOH or increasing concentrations of Ant A (N=3 experiments, One way ANOVA with Dunnett’s multiple comparison test). (G) Ratio of intact MTMR2 band intensity normalized to total (intact + degraded) band intensities from *Parkin^-/-^* neurons treated with EtOH or increasing concentrations of Ant A (N=3 experiments, One way ANOVA with Dunnett’s multiple comparison test). (H) Rubicon band intensity normalized to total protein from *Parkin^-/-^* neurons treated with EtOH or increasing concentrations of Ant A (N=3 experiments, One way ANOVA with Dunnett’s multiple comparison test). (I-L) Inhibition of autophagy does not block the degradation of MTMR5, MTMR2, or Rubicon in response to mitochondrial damage. (I) Representative western blots from lysates of WT cortical neuronal lysates treated with vehicle (DMSO) or 500 nM Bafilomycin A (Baf A1) for 1 hr followed by an additional treatment with vehicle (EtOH) or 15 nM Ant A for 2 hrs. (J) MTMR5 band intensity normalized to total protein from WT neurons treated with DMSO/Baf A1 and EtOH/Ant A. (N=4 experiments, One way ANOVA with Sidak’s multiple comparison test). (K) Ratio of intact MTMR2 normalized to total (intact + degraded) band intensities from WT neurons treated with DMSO/Baf A1 and EtOH/Ant A (N=4 experiments, Kruskal-Wallis test). (L) Rubicon band intensity normalized to total protein from WT neurons treated with DMSO/Baf A1 and EtOH/Ant A (N=4 experiments, One way ANOVA with Sidak’s multiple comparison test). All panels: ns-not significant, ** p<0.01, ***p<0.001, ****p<0.0001, error bars indicate S.E.M.

Pink1/Parkin-mediated mitophagy is a primary degradative mechanism of dysfunctional mitochondria in neurons. Thus, we tested whether degradation of these negative regulators is dependent on Pink1/Parkin mediated mitophagy. For this, we treated embryonic cortical neurons from *Parkin^-/-^* mice with increasing concentration of Ant A. As observed for neurons from wild type mouse embryos, we did not see degradation of the three negative regulators in *Parkin^-/-^*neurons treated with either vehicle (EtOH) or a low dose (3 nM) of Ant A. **(Figures 1E-H)**. Again, as seen for wild type neurons, at higher Ant A concentrations (15 nM, 30 nM) we saw the concerted degradation of MTMR5/2 and Rubicon in *Parkin^-/-^* neurons. Collectively these results indicate that the mitochondrial damage induced degradation of these autophagy suppressors is independent of Pink1/Parkin activity.

The two major degradative pathways in eukaryotic cells are autophagy and the proteasomal system. Given the association of these proteins in negatively regulating autophagy, we first asked if blocking this pathway rescued levels of MTMR5, MTMR2 and Rubicon upon induction of mitochondrial stress. To block autophagosome-lysosome fusion, we treated neurons with Bafilomycin A1 (Baf A1) for 1 hr prior to adding either vehicle or Ant A for another 2 hrs in presence of Baf A1. We found that blocking autophagy did not rescue the loss of MTMR5/2 and Rubicon induced by mitochondrial damage **(Figures 1I-L)**. Instead, we observed a ∼2-fold increase in Rubicon levels, but no increase in the levels of MTMR2/5, when neurons were treated with Baf A1 alone as compared to vehicle **(Figures 1I, 1L)** suggesting that Rubicon undergoes constitutive turnover by the lysosome in neurons, potentially via basal autophagy.

Next, we tested if the degradation of these negative regulators is mediated by the proteasome. We treated neurons with MG132, a drug that blocks proteasome activity, for 1 hr and then added vehicle or Ant A along with MG132 for an additional 2 hrs. We found that inhibiting proteasome activity completely rescued the degradation of MTMR5, MTMR2 and Rubicon upon mitochondrial stress **(Figures 2 A-D)**. We also saw a ∼2-fold increase in Rubicon levels when neurons were treated with MG132 only, in the absence of Ant A, as compared to vehicle-treated neurons **(Figures 2A, D)** suggesting that Rubicon levels are tightly regulated in neurons by both autophagy and the proteasomal pathway. Interestingly, when we treated HeLa cells with either Baf A1 or MG132, we saw no increase **(Supplemental S2A, B)** or only a very minor change **(Supplemental S2C, D)** in Rubicon levels, respectively, indicating that neurons, but not HeLa cells, tightly regulate Rubicon levels even under basal conditions.

**Figure 2:**
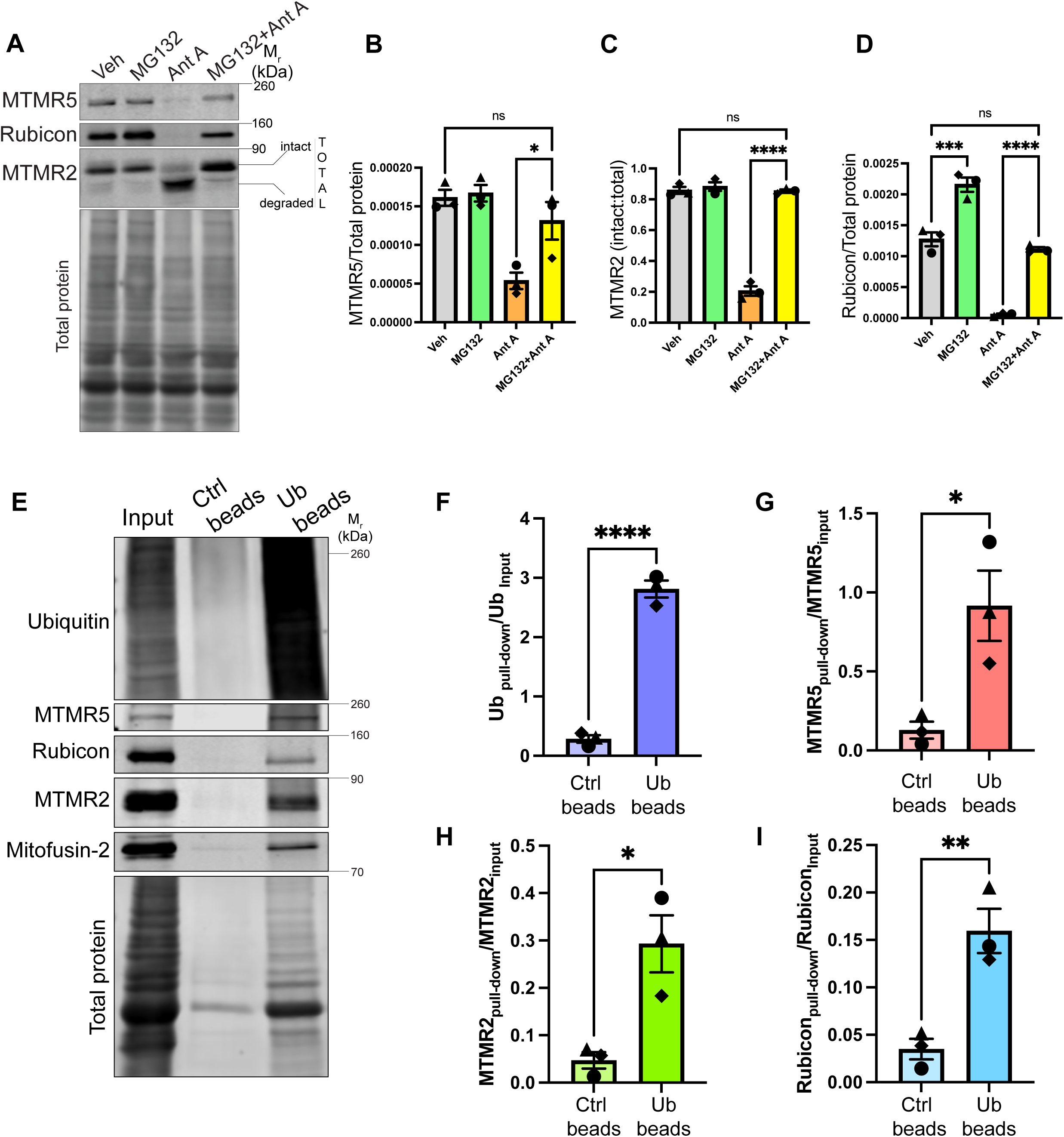
MTMR5, MTMR2 and Rubicon are ubiquitinated and degraded by the proteasome upon acute mitochondrial damage. (A-D) Proteasomal inhibition blocks the concerted degradation of MTMR5, MTMR2, and Rubicon. (A) Representative western blots from lysates of WT murine embryonic cortical neurons treated with vehicle (EtOH) or 10 μM MG132 for 1 hr followed by an additional treatment with vehicle (EtOH) or 15 nM Ant A for 2 hrs. (B) MTMR5 band intensity normalized to total protein from WT neurons treated with EtOH/MG132 and EtOH/Ant A. (N=3 experiments, One way ANOVA with Sidak’s multiple comparison test). (C). Ratio of intact MTMR2 normalized to total (intact + degraded) band intensities from WT neurons treated with EtOH/MG132 and EtOH/Ant A (N=3 experiments, One way ANOVA with Sidak’s multiple comparison test). (D) Rubicon band intensity normalized to total protein from WT neurons treated with EtOH/MG132 and EtOH/Ant A (N=3 experiments, One way ANOVA with Sidak’s multiple comparison test). **(E-I)** MTMR5, MTMR2, and Rubicon are ubiquitinated in response to mitochondrial damage. (E) Representative western blot of ubiquitin enrichment assay from lysates of WT murine embryonic cortical neurons. Neurons were treated with 10 μM MG132 for 1 hr followed by an additional treatment with 15 nM Ant A for another 2 hrs (Ub beads= Ubiquitination affinity beads). (F) Ub enrichment following pull-down with Ub beads or control beads, normalized to intensity in the input lane (N=3 experiments, two-tailed unpaired t-test). (G) MTMR5 enrichment following pull-down with Ub beads or control beads, normalized to intensity in the input lane (N=3 experiments, two-tailed unpaired t-test). (H) MTMR2 enrichment following band intensity after pull-down with Ub beads or control beads, normalized to its intensity in the input lane (N=3 experiments, two-tailed unpaired t-test). (I) Rubicon enrichment after pull-down with Ub beads or control beads, normalized to its intensity in the input lane (N=3 experiments, two-tailed unpaired t-test). All panels: ns-not significant, *p<0.05, ** p<0.01, ***p<0.001, ****p<0.0001, error bars indicate S.E.M.

Proteins that are targeted to the proteasomal machinery for degradation are first conjugated to ubiquitin (Ub) or Ub-like proteins. Thus, we tested if MTMR5/2 and Rubicon are ubiquitinated post mitochondrial damage. For this we treated neurons with 10 µM MG132 for 1 hr to block proteasomal degradation of ubiquitinated proteins and then added 15 nM Ant A for an additional 2 hrs in the presence of MG132 to trigger mitochondrial damage. Post treatment we pulled down Ub-bound proteins from the treated neuronal lysates using a Ub enrichment kit (see Methods). We saw enrichment of Ub in the Ub pull-down fraction when compared to control **(Figures 2 E, F)**. As a positive control, we probed for Mitofusin-2 (MFN-2), as this mitochondrial outer membrane GTPase is ubiquitinated by Parkin as one of the initial steps following induction of PINK1/Parkin-dependent mitophagy^27,65^. Consistent with this, we detected enrichment of MFN-2 in the Ub-pulldown fraction, confirming the onset of mitophagy **(Figures 2E)**. Western blot analysis indicated the presence of all the three negative regulators of autophagy, MTMR5/2 and Rubicon **(Figures 2 E, G-I)** in the Ub-pulldown fraction.

This process of ubiquitination involves a cascade of three sequential enzymatic reactions that include a Ub-activating enzyme (E1), Ub-conjugating enzyme (E2) and a Ub-ligase (E3) ^66^. We then asked if blocking the enzymatic conjugation of Ub to these proteins will prevent their degradation. For this we treated neurons with PYR41, a drug that irreversibly blocks the activity of Ub-E1 enzymes, for 1 hr and then added vehicle or 15 nM Ant A along with PYR41 for additional 2 hrs. We found that blocking the activity of Ub-E1 was sufficient to prevent the degradation of all the three negative regulators of autophagy **(Supplemental S3A)**. Thus, collectively these results indicate that post mitochondrial damage in neurons, the three negative regulators of autophagy are ubiquitinated and undergo dose-dependent degradation by the proteasome, independent of Pink1/Parkin activity.

### The concerted degradation of MTMR5/2 and Rubicon is a specific response to mitochondrial damage in neurons

To test whether the targeted degradation of these three negative regulators is part of a stress response specific to mitochondrial damage, we treated primary cortical neurons with drugs that target different components of the mitochondrial electron transport chain. Oligomycin A (Oligo A) is an inhibitor of mitochondrial ATP synthase (Complex V) thereby blocking oxidative phosphorylation. Carbonyl Cyanide Chlorophenylhydrazone (CCCP) is a mitochondrial uncoupler that interferes with the proton gradient across the inner mitochondrial membrane thereby disrupting mitochondrial membrane potential. Immunoblot analysis indicates that inhibiting mitochondrial ATP synthase by Oligomycin A induced the targeted degradation of MTMR5/2 and Rubicon **(Figure 3 A-D)**. Treatment with the protonophore CCCP at a concentration of 10 μM also elicited a similar response in neurons **(Figure 3 A-D)**. This confirms that either elevating mitochondrial ROS production (by Ant A), inhibiting ATP synthesis (by Oligo A) or disrupting mitochondrial membrane potential (by CCCP) can all trigger the activation of this stress response pathway.

**Figure 3:**
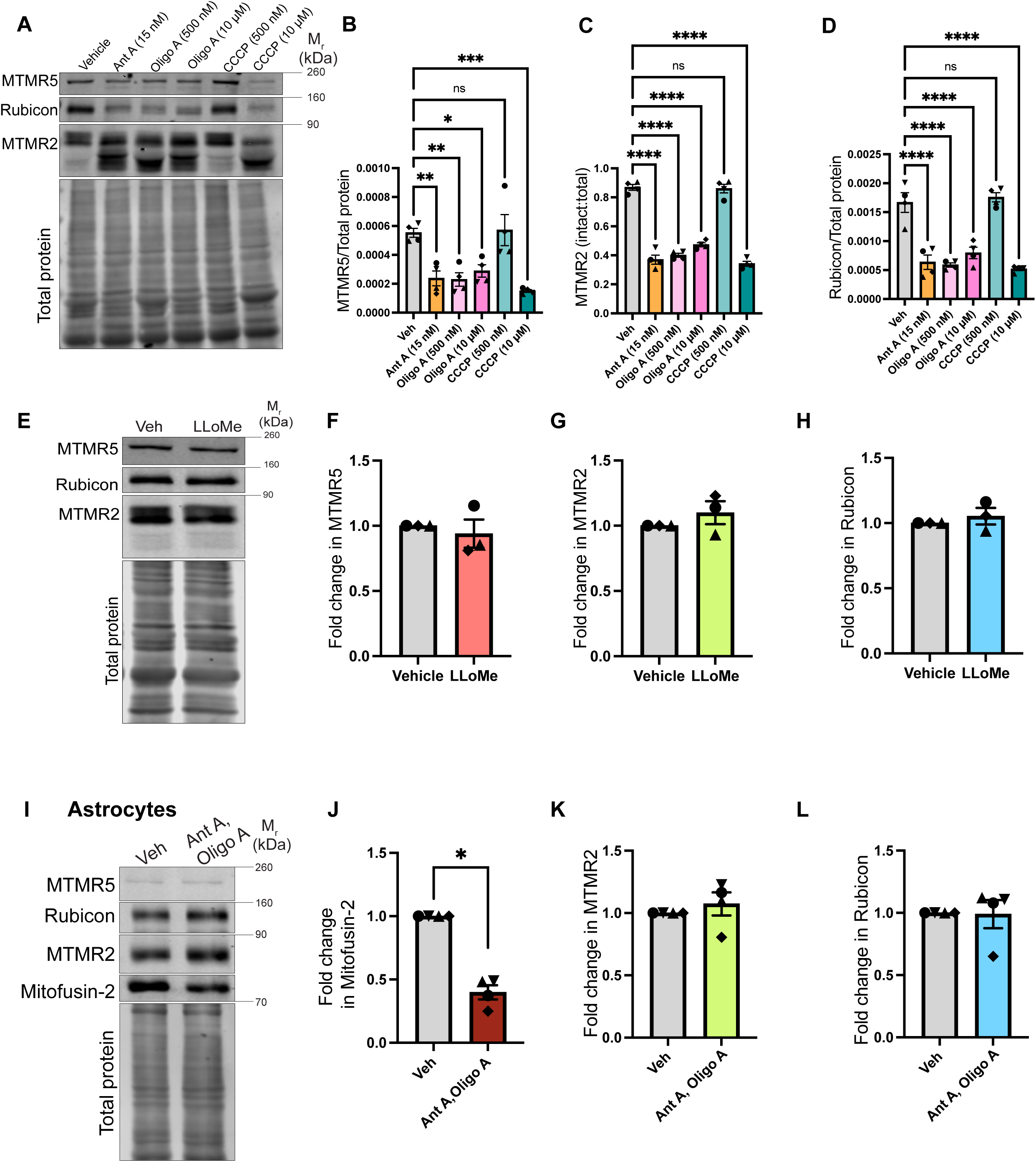
Activation of the MitoSR pathway is specific to mitochondrial damage in neurons and targets negative regulators of autophagy. (A-D) Concerted degradation of MTMR5, MTMR2, and Rubicon is mediated by diverse triggers of mitophagy. (A) Representative western blot from WT cortical neuronal lysates treated with vehicle (EtOH, DMSO), Ant A, Oligo A, or CCCP for 2 hrs (concentration of drugs as indicated in the image). (B) MTMR5 levels normalized to total protein upon treatment of WT neurons with vehicle, Ant A, Oligo A, or CCCP for 2 hrs (N=4 experiments, One way ANOVA with Dunnett’s multiple comparison test). (C) Ratio of intact MTMR2 normalized to total (intact + degraded) upon treatment of WT neurons with vehicle, Ant A, Oligo A, or CCCP for 2 hrs (N=4 experiments, One way ANOVA with Dunnett’s multiple comparison test). (D) Rubicon band intensity normalized to total protein upon treatment of WT cortical neurons with vehicle, Ant A, Oligo A, or CCCP for 2 hrs (N=4 experiments, One way ANOVA with Dunnett’s multiple comparison test). **(E-H)** Lysosomal damage does not trigger the degradation of the negative regulators of autophagy. (E) Representative western blot from lysates of WT murine embryonic cortical neurons treated with vehicle (EtOH) or 1 mM LLoMe for 2 hrs. (F) Fold change in MTMR5 levels upon treatment of WT neurons with LLoMe as compared to with vehicle (N=3 experiments, Mann-Whitney test). (G) Fold change in MTMR2 levels upon treatment of WT neurons with LLoMe as compared to with vehicle (N=3 experiments, Mann-Whitney test). (H) Fold change in Rubicon levels upon treatment of WT neurons with LLoMe as compared to with vehicle (N=3 experiments, Mann-Whitney test). **(I-L)** MitoSR is not triggered by mitochondrial damage in astrocytes. (I) Representative western blot from lysates of WT murine astrocytes treated with vehicle (EtOH, DMSO) or 5 μM Ant A, 10 μM Oligo A for 6 hrs. (J) Fold change in Mitofusin-2 levels upon treatment of WT astrocytes with 5 μM Ant A, 10 μM Oligo A as compared to with vehicle (N=4 experiments, Mann-Whitney test). (K) Fold change in MTMR2 levels upon treatment of WT astrocytes with 5 μM Ant A, 10 μM Oligo A as compared to with vehicle (N=4 experiments, Mann-Whitney test). (L) Fold change in Rubicon levels upon treatment of WT astrocytes with 5 μM Ant A, 10 μM Oligo A as compared to with vehicle (N=4 experiments, Mann-Whitney test). All panels: ns-not significant, *p<0.05, ** p<0.01, ***p<0.001, ****p<0.0001, error bars indicate S.E.M.

We next determined if other forms of organellar damage induced a similar response in neurons. For this, we treated neurons for 2 hrs with 1 mM L-leucyl-L-leucinemethyl ester (LLOMe), a drug that disrupts lysosomal membrane and triggers lysophagy ^67,68^. Immunoblot analysis indicates that although LC3-II levels were elevated following lysosomal damage ^67^ **(Supplemental S3B)**, there was no change in the expression levels of MTMR5/2 or Rubicon **(Figure 3 E-H)**, thereby suggesting that activation of this stress response pathway is specific to mitochondrial damage.

We next checked to see if the degradation of MTMR5/2 and Rubicon was selective, or part of a more generalized loss of autophagy-associated proteins. For this, we looked at levels of VPS34, a component of PI3K complexes that promotes synthesis of PI3P on the growing autophagosome membrane^69,70^. We observed no changes in the levels of VPS34 upon mitochondrial damage **(Supplemental S4 A, B)**. We then tested if other positive regulators of autophagy such as ATG7 and ATG5 are subjected to proteasomal degradation during mitochondrial stress but found their levels to remain unaltered across conditions **(Supplemental S4 C, D)**. We also looked at proteins unrelated to the autophagic machinery such as housekeeping proteins GAPDH, α/β-tubulin and actin. Again, we found that these proteins do not undergo proteasomal degradation upon mitochondrial damage **(Supplemental S4 E-G)**. Thus, we conclude that activation of this pathway is specific for mitochondrial stress and predominantly targets negative regulators of autophagy. Hence, we term this pathway the **Mito**phagic **S**tress **R**esponse (MitoSR) pathway.

Similar to neurons, astrocytes respond robustly to mitochondrial damaging agents to trigger Pink1/Parkin mediated mitophagy^71,72^. In line with this, when we treated primary murine astrocytes with 10 μM Oligo A/5 μM Ant A, we detected a significant loss of MFN-2 in primary astrocytes upon Ant A/Oligo A treatment as compared to vehicle (DMSO/Ethanol) treated cells **(Figure 3 I, J)**, thereby confirming the onset of mitophagy. MTMR5 levels were very low in astrocytes under both conditions, consistent with the identification of MTMR5 as a neuron-enriched protein ^47^. However, MTMR2 **(Figure 3 I, K**) and Rubicon **(Figure 3 I, L)** levels were more robust, and remained comparable across conditions, with no evidence of targeted degradation upon mitochondrial damage. Collectively, this indicates that mitochondrial damage in astrocytes triggers Pink/Parkin mediated mitophagy without activating the MitoSR pathway. Likewise in HeLa cells, upon treatment with Ant A/Oligo A or vehicle (DMSO/Ethanol), we could not detect the degradation of the three negative regulators, although MFN-2 levels were reduced as expected upon Ant A/Oligo A treatment **(Supplemental S5 A-F)**. Taken together, these results suggest that the degradation of the negative regulators of mitophagy, MTMR5/2 and Rubicon, by the MitoSR pathway represents a neuron-specific response to acute levels of mitochondrial damage.

### Rubicon localizes to somal lysosomes in neurons

We then questioned how the targets of the MitoSR pathway i.e. MTMR5, MTMR2 and Rubicon affect neuronal autophagy under otherwise basal conditions. MTMR5 and MTMR2 act together to repress the early steps of neuronal autophagy by inhibiting PI3P formation on autophagosomes ^47^, and thus degradation of these proteins would be predicted to enhance autophagosome biogenesis. However, the role of Rubicon in regulating autophagy in neurons, if any, remains unknown. We probed for endogenous Rubicon in neurons and found a ∼3.5 fold enrichment when compared to HeLa cells **(Figure 4A, B)**. To examine the subcellular localization of Rubicon, we transfected neurons with an EGFP-tagged construct of human Rubicon and performed live imaging. We found Rubicon to form ring-like punctate structures that were predominantly distributed within the neuronal soma **(Figure 4C)**; punctate structures were also observed in both dendrites **(Figure 4D)** and axons **(Figure 4E)**, but less frequently. We co-transfected neurons with an autophagosome marker (mCherry-LC3) and the lysosomal marker (LAMP1-Halo) along with GFP-Rubicon. Almost 60% of GFP-Rubicon puncta within the soma co-localized with LAMP1-Halo, while only 10% of GFP-Rubicon puncta colocalized with mCherry-LC3 **(Figure 4F, G)**. Thus, in neurons, as well as the non-neuronal lines A549 and HeLa cells^52,53^, Rubicon is localized to lysosomes and late endosomes.

**Figure 4:**
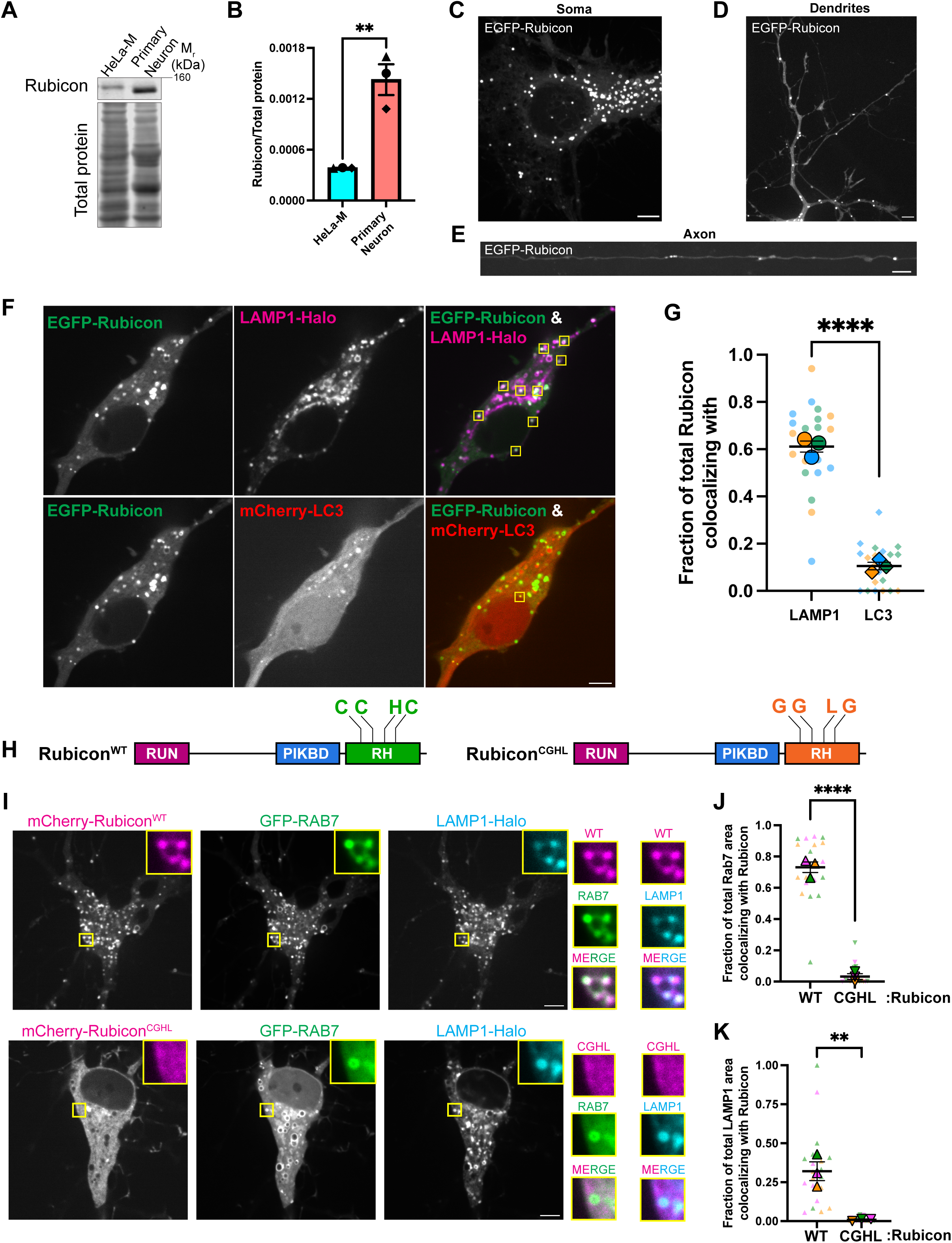
Rubicon localization to lysosomes in neurons is Rab7-dependent. (A, B) Rubicon expression is enhanced in neurons. (A) Representative western blot from lysates of HeLa cells or WT murine embryonic cortical neurons and probed for Rubicon. (B) Rubicon band intensity normalized to total protein in HeLa cells or WT neurons (N=3 experiments, two-tailed unpaired t-test). **(C-E)** Rubicon localizes to organelles distributed throughout the soma, dendrites, and axons of primary cortical neurons. (C) Representative max projection of the soma of a WT cortical neuron transfected with EGFP-Rubicon. (D) Representative max projection of the dendrites of a WT cortical neuron transfected with EGFP-Rubicon. (E) Representative max projection of an axon of a WT cortical neuron transfected with EGFP-Rubicon. **(F, G)** Rubicon colocalizes with the late endosome/lysosome marker LAMP1-Halo but not with the autophagosome marker mCherry-LC3. (F) Single z-plane confocal images of the soma of a WT cortical neuron transfected with mCherry-LC3, LAMP1-Halo and EGFP-Rubicon. Yellow boxes indicate EGFP-Rubicon colocalizing with either LAMP1-Halo or mCherry-LC3. (G) Fraction of the number of EGFP-Rubicon puncta in the soma of each neuron colocalizing with LAMP1-Halo or mCherry-LC3 (N=3 experiments, two-tailed unpaired t-test). **(H-K)** Rubicon localization to lysosomes is Rab7-dependent. (H) Schematics of the domain organization of Rubicon^WT^ and Rubicon^CGHL^ with the RUN, PI3K-binding domain (PIKBD) and Rubicon homology (RH) domains annotated. (I) Single z-plane confocal images of somas of neurons transfected with GFP-Rab7, LAMP1-Halo and either mCherry-Rubicon^WT^ or mCherry-Rubicon^CGHL^. Yellow boxes indicate inset regions. (J) Fraction of total GFP-Rab7 area in each neuronal soma colocalizing with mCherry-Rubicon^WT^ and mCherry-Rubicon^CGHL^ (N=3 experiments, two-tailed unpaired t-test). (K) Fraction of total Halo-LAMP1 area in each neuronal soma colocalizing with mCherry-Rubicon^WT^ and mCherry-Rubicon^CGHL^ (N=3 experiments, two-tailed unpaired t-test). All panels: ** p<0.01, ****p<0.0001, error bars indicate S.E.M., scale bars= 5μm.

Rubicon binds to Rab7 via its C-terminal RH domain **(Figure 4H)**, and this interaction negatively impacts endocytic trafficking and autophagosome maturation in non-neuronal cells such as HEK293 or HeLa^58–60^. Within this RH domain, mutations of three conserved cysteine residues to glycine (*C912G*, *C915G*, *C923G*) and the histidine residue to leucine (*H920L*) (Rubicon^CGHL^) result in a complete loss of Rubicon and Rab7 interaction ^58,68^. To test if this domain determines Rubicon’s localization in neurons, we introduced the same mutations in the full-length Rubicon protein **(Figure 4H)** and tracked its localization in neurons. We co-expressed either mCherry-Rubicon^WT^ or mCherry-Rubicon^CGHL^ in neurons along with GFP-Rab7 and LAMP1-Halo. We found that while Rubicon^WT^ showed a strong membrane association and co-localization with both Rab7 and LAMP1, Rubicon^CGHL^ remained cytosolic and no longer co-localized with either Rab7 or LAMP1 **(Figure 4 I-K)**. This observation suggests that the RH domain of Rubicon is critical for the association of Rubicon with lysosomes in neurons.

### Rubicon represses lysosomal function in neurons

Having determined the basis of the stark localization of Rubicon in the neurons, we next examined if Rubicon plays a role in regulating neuronal autophagy under basal conditions. Within the neuronal soma, the majority of LC3-positive organelles are usually autolysosomes that are also positive for LAMP1 ^46,74^. Interestingly, however, we found that the majority of lysosomes (marked by LAMP1-Halo) that are associated with EGFP-Rubicon do not colocalize with autophagosomes (marked by mCherry-LC3) **(Figure 5 A, B)**. Hence, we questioned if Rubicon blocks autophagosome-lysosome fusion in neurons. For this, we transfected neurons with either EGFP or EGFP-Rubicon along with mCherry-LC3 and LAMP1-Halo. We measured autophagosome numbers per neuronal soma and found no change in the number of mCherry-LC3 puncta per cell upon overexpression of Rubicon **(Figure 5 C, D)**. However, when we measured the colocalization between mCherry-LC3 and LAMP1-Halo as a read out of autolysosome formation, we found overexpression of EGFP-Rubicon led to a ∼50% decrease in mCherry-LC3 and LAMP1-Halo colocalizing puncta **(Figure 5 C, E)**. To determine whether this decrease in autolysosome numbers was due to reduced autophagosome-lysosome fusion or due to an inhibition of autophagosome biogenesis, we quantified the ratio of LAMP1^+^LC3^+^ autolysosomes to LC3^+^ autophagosome in EGFP or EGFP-Rubicon overexpressing neurons. In EGFP-overexpressing neurons, we found almost ∼70% autophagosomes colocalized with lysosomes; this number decreased to ∼34% in neurons overexpressing GFP-Rubicon, indicating a ∼50% drop in autophagosome-lysosome fusion **(Figure 5 C, F)**. Therefore, we demonstrate that overexpression of Rubicon reduced autophagy flux primarily by blocking autolysosome formation.

**Figure 5:**
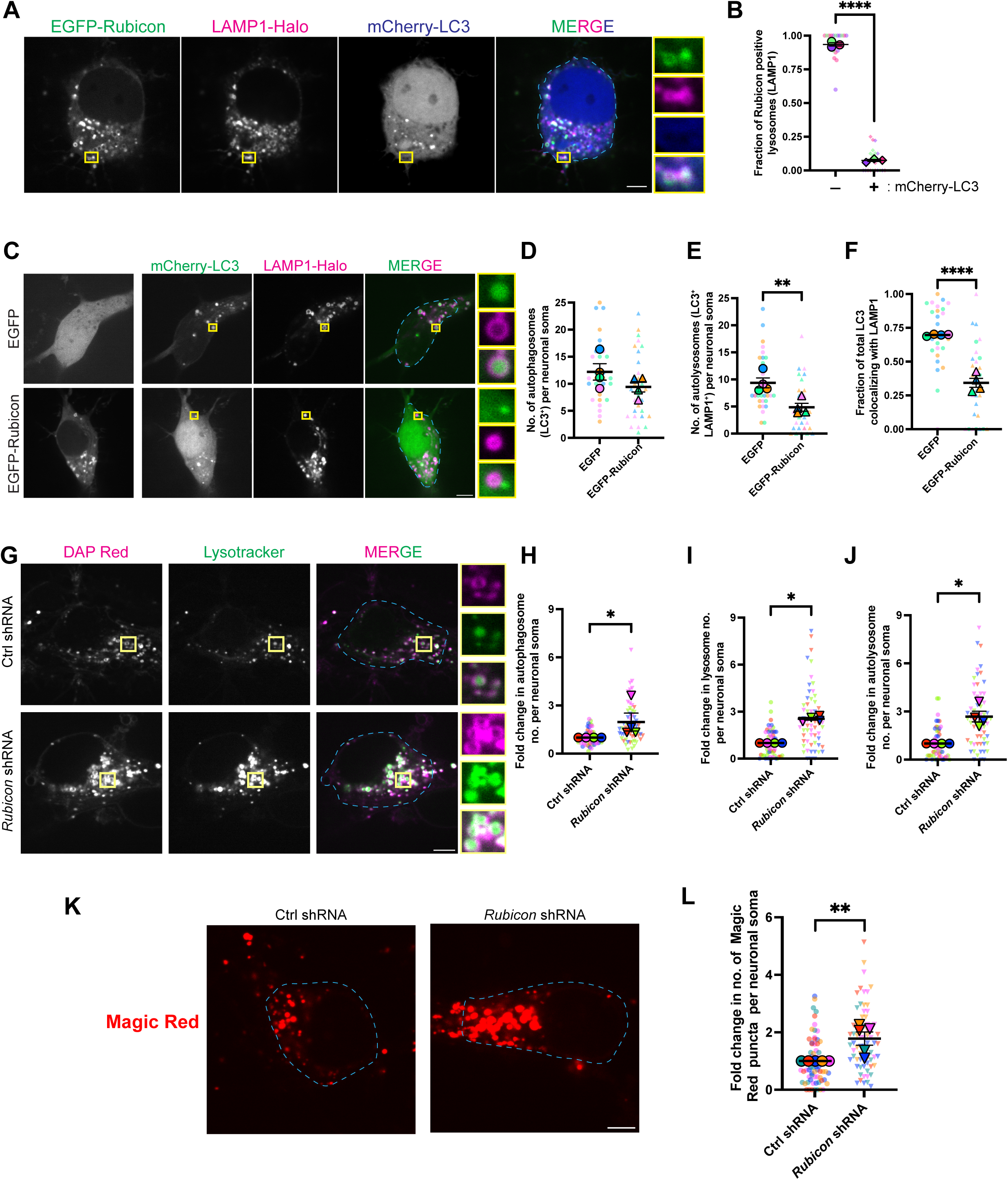
Rubicon inhibits lysosomal function and autophagosomal maturation. (A, B) Rubicon is associated with lysosomes but not autolysosomes. (A) Single z-plane confocal images of a neuron transfected with EGFP-Rubicon, LAMP1-Halo and mCherry-LC3. Outline of the neuronal soma for quantification is indicated using dashed lines in cyan. Yellow boxes indicate inset regions. (B) Fraction of lysosomes (LAMP1-Halo) colocalizing with EGFP-Rubicon in the soma that either colocalize with mCherry-LC3 or not (N= 3 experiments, two-tailed unpaired t-test). **(C-F)** Overexpression of Rubicon inhibits autophagosome maturation. (C) Single z-plane confocal images of neurons transfected with mCherry-LC3, LAMP1-Halo and either EGFP or EGFP-Rubicon. Outline of the neuronal soma for quantification is indicated using dashed lines in cyan. Yellow boxes indicate inset regions. (D) Number of autophagosomes (marked by mCherry-LC3 punctae) per neuronal soma either expressing EGFP or EGFP-Rubicon (N= 4 experiments, two-tailed unpaired t-test). (E) Number of autolysosomes (marked by colocalizing mCherry-LC3 and LAMP1-Halo punctae) per neuronal soma either expressing EGFP or EGFP-Rubicon (N= 4 experiments, two-tailed unpaired t-test). (F) Fraction of autolysosomes (LAMP^+^LC3^+^) to autophagosomes (LC3^+^) per neuronal soma either expressing EGFP or EGFP-Rubicon (N= 4 experiments, two-tailed unpaired t-test). **(G-J)** Rubicon knockdown increases the number of lysosomes, autophagosomes, and autolyososomes in cortical neurons. (G) Single z-plane confocal images of neurons nucleofected with control or *Rubicon* shRNA and treated with DAP Red and Lysotracker. Outline of the neuronal soma for quantification is indicated using dashed lines in cyan. Yellow boxes indicate inset regions. (H) Fold change in autophagosome numbers (marked by DAP Red punctae) per neuronal soma in control vs Rubicon knockdown neurons (N=4 experiments, Mann-Whitney test). (I) Fold change in lysosome numbers (marked by Lysotracker punctae) per neuronal soma in control vs Rubicon knockdown neurons (N=4 experiments, Mann-Whitney test). (J) Fold change in autolysosome numbers (marked by colocalizing DAP Red and Lysotracker punctae) per neuronal soma in control vs Rubicon knockdown neurons (N=4 experiments, Mann-Whitney test). **(K, L)** Rubicon knockdown increases the number of proteolytically active lysosomes. (K) Single z-plane confocal images of neurons nucleofected with control or *Rubicon* shRNA and treated with Magic Red dye. Outline of the neuronal soma for quantification is indicated using dashed lines in cyan. Yellow boxes indicate inset regions. (L) Fold change in the number of Magic Red punctae per neuronal soma in control vs Rubicon knockdown neurons (N=5 experiments, Mann-Whitney test). All panels: *p<0.05, ** p<0.01, ****p<0.0001, error bars indicate S.E.M., scale bars = 5μm.

We then determined the molecular mechanism by which Rubicon blocks autophagosome-lysosome fusion in neurons. For this, we knocked down endogenous Rubicon by nucleofecting neurons with a targeted shRNA plasmid. After 6-7 days of knockdown **(Supplemental S6 A, B)**, we labeled autophagosomes using the autophagy dye DAP Red, which incorporates into the autophagosome membrane and fluoresces in a hydrophobic environment^75–77^, then labeled lysosomes using Lysotracker which fluoresces in the acidic environment of the lysosome. Compared to control neurons, Rubicon depletion resulted in a ∼2 fold increase in the number of autophagosomes marked by DAP Red **(Figure 5 G, H)**. We saw an even greater increase (∼2.5 fold) in the number of acidic lysosomes marked by Lysotracker in Rubicon knockdown neurons **(Figure 5 G, I)**, indicating that Rubicon impairs lysosomal acidification. As a read-out of the number of autolysosomes formed, we quantified the number of autophagosomes colocalizing with Lysotracker, and found a ∼2.7-fold increase in the number of autolysosomes per soma when Rubicon was depleted from neurons **(Figure 5 G, J)**. In concert with our observations upon Rubicon overexpression, these data indicate that Rubicon effectively blocks autophagosome-lysosome fusion, and that degradation of Rubicon enhances autophagy flux. We hypothesized that the mechanism by which Rubicon directly affects autophagosome-lysosome fusion is by localizing to lysosomes and modulating organelle function. To test this, we knocked down Rubicon and measured the abundance of functional active lysosomes, using the Magic Red dye which fluoresces upon hydrolysis by the lysosomal protease Cathepsin B. We found a ∼1.8-fold increase in the number of Magic Red punctae **(Figure 5 K, L)** upon Rubicon knockdown, demonstrating a clear role for Rubicon in regulating lysosomal activity. Thus, we identify that in neurons, Rubicon halts autophagosome-lysosome fusion via regulating lysosomal function.

### Depletion of negative regulators promotes mitophagy flux in neurons

Previous work has shown that MTMR5-MTMR2 suppress autophagosome formation in neurons ^47^, and we find that Rubicon negatively regulates lysosomal function to modulate autophagosome-lysosome fusion. Thus, during acute mitochondrial stress when the MitoSR pathway is activated, the concerted degradation of MTMR5, MTMR2 and Rubicon upregulates autophagic flux to promote the efficient turnover of damaged mitochondria. Hence, we asked whether depletion of these negative regulators would be sufficient to promote more efficient mitophagic flux in neurons undergoing mild oxidative damage; levels that are below the threshold to trigger the MitoSR pathway **(Figure 1 A-3nM Ant A)**. From a disease perspective, this might represent a powerful therapeutic approach, as it is thought that most neurological disorders where autophagy is compromised are chronic in nature with the neurons undergoing mild amounts of stress over time; levels that would be insufficient to activate the MitoSR pathway. For this, we nucleofected neurons with shRNA plasmids against *Rubicon* and the active PI3P phosphatase *Mtmr2* at DIV=0. After 6-7 days of knockdown **(Supplemental S6 C-E)**, we performed a mitophagy flux assay **(Figure 6A)**. In this assay we labeled neurons with the mitophagy dye DMP Red, which primarily labels acidified mitochondria, indicating that they are engulfed within an acidified autolysosome ^78–80^. To induce mild mitochondrial damage, we treated neurons with 3 nM Ant A ^35^ for 1 hr and chased for another 2 hrs to probe for change in mitophagy flux. As an additional test, we used Lysotracker to label acidic lysosomes and checked for colocalization between the mitophagy dye and Lysotracker to quantify the number of mitophagolysosomes. Compared to neurons treated with a control shRNA, combinatorial knockdown of MTMR2 and Rubicon resulted in a significant increase in the number of mitophagolysosomes **(Figure 6B, C)**. The combined depletion of MTMR2 and Rubicon resulted in a ∼3 -fold increase in mitophagy flux in neurons undergoing very low levels of mitochondrial stress. This observation suggests that the targeted depletion of the negative regulators of autophagy can result in enhanced mitochondrial turnover under more physiological levels of stress.

**Figure 6:**
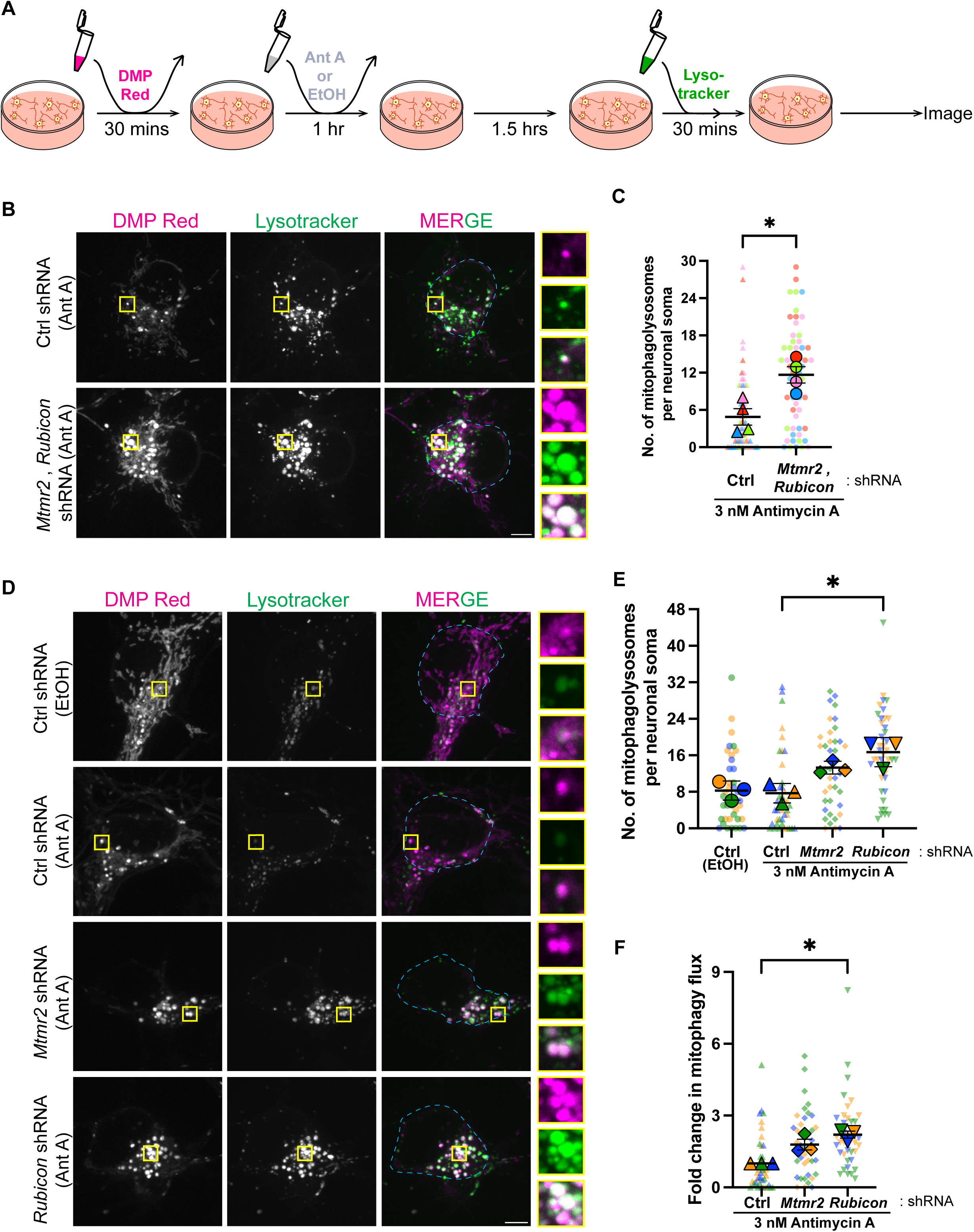
Depletion of MTMR2 and Rubicon promotes autophagic clearance of damaged mitochondria. (A) Schematic depicting the steps of the mitophagy assay using DMP Red and Lysotracker to label acidified mitochondria and lysosomes, respectively. **(B, C)** Depletion of MTMR2 and Rubicon increases the number of mitochondria engulfed within autophagosomes. (B) Representative max projections of neurons nucleofected with Ctrl or *Mtmr2* and *Rubicon* shRNA and assayed for mitophagy using the protocol represented in Fig:6A. Outline of the neuronal soma for quantification is indicated using dashed lines in cyan. Yellow boxes indicate inset regions. (C) Number of mitophagolysosomes per soma (marked by colocalizing DMP Red and Lysotracker punctae) of neurons nucleofected with Ctrl or *Mtmr2* and *Rubicon* shRNA and assayed for mitophagy using the protocol represented in Fig:6A (N=4 experiments, two-tailed unpaired t-test). **(D-F)** Depletion of Rubicon is sufficient to significantly enhance mitophagic flux. (D) Representative max projections of neurons nucleofected with Ctrl, *Mtmr2* or *Rubicon* shRNA and assayed for mitophagy using the protocol represented in Fig:6A. Outline of the neuronal soma for quantification is indicated using dashed lines in cyan. Yellow boxes indicate inset regions. (E) Number of mitophagolysosomes per soma (marked by colocalizing DMP Red and Lysotracker punctae) of neurons nucleofected with Ctrl, *Mtmr2* or *Rubicon* shRNA and assayed for mitophagy using the protocol represented in Fig:6A (N=3 experiments, Kruskal-Wallis test). (F) Fold change in mitophagy flux per soma of neurons nucleofected with Ctrl, *Mtmr2* or *Rubicon* shRNA and assayed for mitophagy using the protocol represented in Fig:6A (N=3 experiments, Kruskal-Wallis test). All panels: *p<0.05, error bars indicate S.E.M, scale bars = 5μm.

Since MTMR2 affects early steps of autophagosome biogenesis, and Rubicon predominantly the later steps, we asked if they act independently or in concert to repress mitophagy. To test this, we individually knocked down MTMR2 **(Supplemental S6 F, G)** and Rubicon **(Supplemental S6 A, B)** in neurons and tested the impact on mitophagy flux upon mild mitochondrial damage (3 nM Ant A). We found that depletion of MTMR2 resulted in a modest increase in the number of mitophagolysosomes as compared to control **(Figure 6 D, E)**. However, in neurons where Rubicon was knocked down, we observed a significant increase in the number of mitophagolysosomes **(Figure 6 D, E)**. When we quantified the change in mitophagy flux we saw that knockdown of Rubicon resulted in ∼2-fold increase in mitophagic flux **(Figure 6F)**. It is also interesting to note that the number of mitophagosomes does not change significantly between vehicle-treated and Ant A-treated control neurons with wild type levels of Rubicon **(Figure 6 D, E)**. This observation reinforces our finding that levels of negative regulators, especially Rubicon, are unchanged under mild oxidative stress **(Figure 1 A-** vehicle vs 3 nM) and these levels are sufficient to inhibit mitochondrial turnover. Additionally, our results demonstrate that knocking down Rubicon had a greater impact on mitophagy flux than MTMR2 depletion **(Figure 6F)**, potentially due to the fact that MTMR2 acts early in the autophagy pathway while Rubicon predominantly affects the penultimate step which is blocking autophagosome-lysosome fusion.

## DISCUSSION

Mitochondrial dysfunction can have devastating effects on neuronal health, and has been implicated in the pathogenesis of several neurological disorders including PD or ALS ^81^. The post-mitotic nature of neurons makes it essential that these cells either repair or degrade damaged mitochondria in order to prevent the release of ROS, Ca^2+^, and/or mitochondrial DNA^81–83^. Here, we report a novel stress response pathway that is triggered in neurons by high levels of mitochondrial damage. This pathway, which we term the Mitophagic Stress Response, or MitoSR pathway, operates in parallel to the classical Pink1/Parkin-mediated mitophagy process **(Figure 7)**. Both mechanisms are induced upon mitochondrial stress in neurons, and together these mechanisms lead to effective engulfment and degradation of damaged mitochondria. Activation of MitoSR triggers the concerted degradation of MTMR5, MTMR2 and Rubicon by the ubiquitin-proteosome system; these proteins actively suppress autophagic flux in distinct but via complementary mechanisms. Further, we find that targeted depletion of MTMR2 and Rubicon is sufficient to significantly stimulate neuronal mitophagy under mild mitochondrial stress that are below the threshold activating the MitoSR pathway.

**Figure 7:**
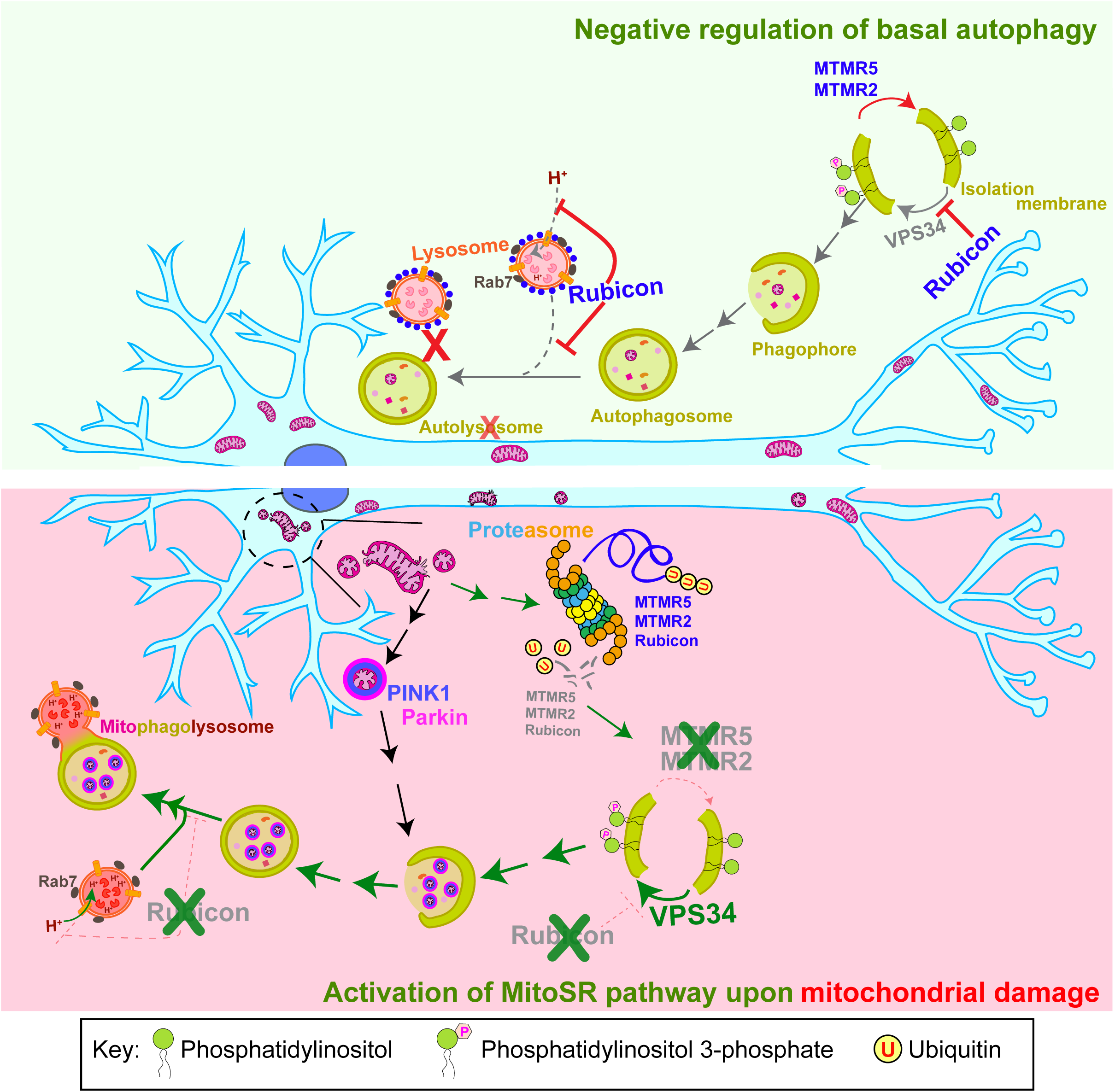
**MTMR5/2 and Rubicon suppress neuronal autophagy under basal conditions, while acute mitochondrial stress induces the selective degradation of these negative regulators via the MitoSR pathway**. The upper panel depicts the roles of MTMR5/2 and Rubicon in repressing neuronal autophagy under basal conditions. MTMR5/2 activity hydrolyzes PI3P ito PI, inhibiting early steps of autophagosome biogenesis ^47^. Rubicon blocks lysosomal acidification and autophagosome/lysosome fusion, thereby impacting latter steps of autophagy. The lower panel depicts induction of the MitoSR pathway, which acts in parallel to the Pink1/Parkin pathway for mitophagy initiation in response to mitochondrial damage in neurons. Activation of MitoSR induces the ubiquitination of MTMR5, MTMR2 and Rubicon and the targeting of these negative regulators to the proteasome for degradation. This concerted degradation facilitates increased autophagic flux, promoting mitochondrial turnover.

Our results show that the degradation of negative regulators of autophagy by MitoSR is brought about by the proteasomal machinery, and not by autophagosomes. The proteasomal machinery has previously been reported to regulate levels of several autophagic proteins (WIPI2^84^, ULK1^85^, ATG16L^86^) under basal conditions. Independent of this, mitochondrial damage also triggers proteasomal degradation of several mitochondrial outer membrane proteins ^87,88^. However, the activation of the MitoSR pathway represents a coordinated signaling response between mitochondrial stress, the ubiquitin-proteasomal machinery, and autophagy. It is indeed striking that the MitoSR pathway is specific for mitochondrial stress, as when we induced lysosomal damage in neurons using LLoMe^67^, we did not detect degradation of any of the negative regulators. Additionally, we observed that the MitoSR pathway does not get activated in primary astrocytes or HeLa cells, indicating that activation of this pathway may be a neuron-specific response to high levels of mitochondrial stress. We speculate that the specificity of this pathway arises due to the presence of neuron-enriched E3 ligase(s) or to intrinsic mitochondrial proteins that are activated during mitochondrial damage; further work will be required to differentiate between these possibilities.

Along with the MTMR5-MTMR2 complex which has previously been reported to suppress autophagosome biogenesis ^47^, here we identify Rubicon as a critical negative regulator of basal neuronal autophagy **(Figure 7)**. Rubicon is abundantly expressed in neurons and localizes to somal LAMP1-positive endosomes or lysosomes. In accordance with previous key studies on Rubicon in non-neuronal systems ^52,53,57–60^, we find that in neurons, Rubicon binds to Rab7 and is recruited to lysosomal membranes via its N-terminal RH domain. Contrary to previous reports, however, we find that in neurons Rubicon has a more dramatic effect on blocking autophagosome-lysosome fusion. When Rubicon is depleted, there is a significant enrichment in the number of acidic lysosomes and an even greater increase in the number of autolysosomes. In accordance with this, we observe increased lysosomal function (as measured by Cathepsin B activity) when Rubicon is knocked down in neurons. Thus, in neurons, Rubicon associates with lysosomes **(Figure 4F, G)** and modulates lysosomal acidification **(Figure 5 G, I)** and thereby their hydrolytic activity **(Figure 5 K, L)**. This inhibition directly impacts the latter steps of autophagy, primarily affecting autophagosome-lysosome fusion **(Figure 5C, F**; **Figure 5 G, J)**. Thus, MTMR5/2 represses the early steps of autophagy ^47^, while Rubicon predominantly regulating the later steps, both of which are critical regulatory nodes in the context of neurological disorders where autophagy is compromised ^89^.

Previous work from our lab has shown that fusion of mitophagosomes with lysosomes is slow, and in fact is the rate limiting step in neuronal mitophagy ^35^. Here, we provide a mechanistic basis for this delay in fusion. At low levels of damage (3 nM), Rubicon is not degraded **(Figure 1A, D)** and given its role in blocking autophagosome-lysosome fusion **(Figure 5C, F)**, it hinders mitophagosome acidification by lysosomes. However, when the MitoSR pathway is activated at higher levels of damage, the degradation of Rubicon facilitates the fusion of autophagosomes carrying damaged mitochondria with lysosomes. Consistent with this hypothesis, when we deplete Rubicon levels in neurons and induce mild mitochondrial damage, we observe a twofold increase in mitophagy flux. Additionally, we also saw a modest increase in mitophagy flux upon depleting MTMR2. As a part of the MitoSR pathway, knockdown of MTMR2 will upregulate autophagosome formation, but because Rubicon controls the fusion of these autophagosomes with lysosomes downstream, it acts as the limiting factor in regulating flux.

Our work opens interesting avenues for future investigation. Why do neurons require such tight negative regulation of autophagy? What happens if autophagy is upregulated in neurons either acutely or chronically? An interesting study in proximal tubular epithelial cells showed that in the absence of Rubicon, lysosomal function is overwhelmed ^90^. There is excess transfer of phospholipids from different organellar membranes to lysosomes via autophagy, leading to formation of expanded hyperactive lysosomes. Our work suggests that in neurons, Rubicon also acts as a critical check point to regulate lysosomal function. By regulating lysosomal physiology, Rubicon may control basal autophagy flux to prevent dysregulated breakdown of cellular components by autolysosomes. This raises an interesting question - what impact does Rubicon’s role in blocking autophagosome-lysosome fusion have on secretory autophagy? Inhibiting autophagosome maturation or lysosomal function pharmaceutically has been shown to trigger the release of extracellular vesicles harboring autophagic receptors such as p62, NDP52, OPTN, NBR1 along with LC3-II ^91,92^, in a process described as ‘secretory autophagy during lysosome inhibition (SALI)’. We predict that in neurons, Rubicon maintains a fine balance between conventional autophagy and SALI by modulating lysosomal physiology and autolysosome numbers.

In summary, here we identify a novel stress response pathway, the MitoSR pathway, which is activated under intense mitochondrial damage to modulate levels of the negative regulators of neuronal autophagy. We uncover a mechanistic role of Rubicon in repressing neuronal autophagy by regulating lysosomal function. Further, we show that under mild mitochondrial stress, the targeted depletion of these negative regulators can promote autophagic turnover of mitochondria. This finding may be of particular importance during chronic neurodegenerative disease where mitochondrial stress may be mild but continue to build over time. We thus propose that in degenerating neurons of patients with PD or ALS where mitophagy is compromised, targeting negative regulators of autophagy may enhance mitochondrial turnover and prevent the pathological accumulation of dysfunctional mitochondria that is characteristic of these diseases.

## MATERIALS AND METHODS

### Chemicals and Reagents

The following chemicals were used: Ethanol (Decan Labs, #2716), Dimethyl sulfoxide (DMSO, Sigma Aldrich, #D2650), Antimycin A (Sigma Aldrich, #A8674), Bafilomycin A1 (Sigma, Aldrich, #SML1661), MG132 (Selleckchem, #S2619), Oligomycin A (Sigma Aldrich, #75351), CCCP (Sigma Aldrich #C2759), LLoMe (Cayman Chemicals #16008), PYR 41 (Tocris Bioscience #2978). The following reagents were used for *in vitro* cultures: Poly-L-Lysine (Sigma Aldrich #P1274), Poly-D-Lysine (Gibco #A3890401), Horse Serum (Gibco #16050122), Fetal Bovine Serum (Sigma Aldrich #F2442), Sodium Pyruvate (Gibco #11360070), D-(+)-Glucose solution (Sigma Aldrich #G8769), Sodium Chloride (NaCl, Fisher BioReagents #BP358-1), Minimal Essential Medium (MEM, Gibco #11095072), GlutaMAX (Gibco # 35050061), Penicillin-Streptomycin (10000 U/mL, Gibco #15140122), Neurobasal Medium (Gibco # 21103049), B27 Supplement-serum free (Gibco #17504044), Hibernate® E Low Fluorescence (TransnetyxYX Tissue *#*HELF-500 ml), AraC (Sigma Aldrich, #C6645), Dulbecco’s Modified Eagle’s Medium (DMEM, Corning, #10-017-CM), DMEM-high glucose & pyruvate (Gibco #11995073).

### Experimental model and subject details

Mouse lines used in this study include wild type [C57BL/6J (RRID: IMSR_JAX:000664)] and Parkin^-/-^ [B6.129S4-Prkntm1Shn/J (RRID: IMSR_JAX:006582)]. Cortices were dissected out from mouse embryos at day 15.5 in accordance with protocols approved by the Institutional Animal Care and Use Committee at the University of Pennsylvania. Neurons were isolated by digestion with 0.25% Trypsin followed by trituration. The neuronal suspension was then plated in attachment media (MEM and supplements including 10% horse serum, 1 mM sodium pyruvate, 33 mM D-glucose and 37.5 mM NaCl) on poly-L-Lysine coated plates. Post attachment (5-6 hrs), the media was replaced with maintenance media (MM) [Neurobasal and supplements including with 33 mM D-glucose, 2mM GlutaMax, 37.5 mM NaCl, Penicillin (100units/mL)/Streptomycin (100 mg/mL), and 2% B27]. On the following day 5 µM of AraC was added to restrict the growth of non-neuronal cells such as glia. For experiments involving knockdown with shRNA plasmid, 6-7 days *in vitro* (DIV) neurons were used. Other experiments were performed with DIV 8-10 neurons.

Primary murine astrocytes were isolated based on previously published protocols^72,93^.

HeLa-M cells (referred to as HeLa in the main text, a gift from Andrew Peden, Cambridge Institute for Medical Research, UK) were authenticated and checked for mycoplasma contamination by STR profiling and MycoAlert detection kit (Lonza, LT07) respectively. Cells were grown in Dulbecco’s Modified Eagle’s Medium and supplemented with 1% GlutaMax and 10% Fetal Bovine Serum and 1% GlutaMax.

Neurons, astrocytes, and HeLa cells were grown in separate incubators at 37^0^C with 5% CO_2_.

### Transfection/Nucleofection

Mouse embryonic cortical neurons were transfected using Lipofectamine 2000 (ThermoFisher Scientific Cat#11668019). A total of 1.5 µg DNA was mixed with 2 µl Lipofectamine and incubated for 30 mins at room temperature (RT) to form DNA-lipid complexes. The mixture was then added to neurons in fresh MM and incubated for 90 min post which the media was replaced with 1:1 conditioned: fresh MM. Neurons were transfected for 18-24 hrs prior to imaging. The following plasmids were used for transfections: EGFP-Rubicon (RRID: Addgene 221657), LAMP1-Halo (RRID: Addgene 221655), mCherry-LC3 (RRID: Addgene 221656), mCherry-Rubicon^WT^ (RRID: Addgene 221658), mCherry-Rubicon^CGHL^ (RRID: Addgene 221659) and EGFP-Rab7A (RRID: Addgene 28047), EGFP-N1 (RRID: Addgene 6085-1). HeLa cells were transfected with 1.5 µg of untagged human Parkin (RRID: Addgene 187897) using Lipofectamine 2000 for 18-24 hrs prior to treatment.

On the day of dissection, post trypsinization, embryonic neurons in suspension were nucleofected with shRNA plasmid. Nucleofections were carried out in an Amaxa nucleofector II device from Lonza according to the manufacturer’s protocol. For each 10^6^ neurons, 3 µg of shRNA plasmid was used for nucleofecting. The plasmids used were: Non-Target shRNA Control Plasmid DNA (Sigma Aldrich; SHC016), TRC mouse *Rubicon* shRNA (Sigma Aldrich; TRCN0000283271), TRC mouse *Mtmr2* shRNA (Sigma Aldrich; TRCN0000030094)

### Western blotting

Samples were lysed in RIPA buffer (50 mM Tris-HCl, 150 mM NaCl, 0.1% Triton X-100, 0.5% sodium deoxycholate and 0.1% SDS, pH=7.4) supplemented with Halt Protease and phosphatase inhibitor (ThermoFisher Scientific Cat#78442) at 4^0^C for 30 mins. Samples were then centrifuged at 17000X g and the supernatant was collected. Protein concentration from the supernatant was measured using Pierce BCA Protein Assay Kit (ThermoFisher Scientific, Cat#23225). Samples were then denatured at 95^0^C and resolved on an SDS-PAGE gel. The resolved proteins on the gel were then transferred to Immobilon-FL PVDF membranes (Millipore) at 100V for 1 hr 12 mins. Post transfer, membranes were dried for 1 hr at RT and then rehydrated using methanol. The membranes were then stained with Revert™ 700 Total Protein Stain (Licor Cat#926-11021) for 5 mins and washed twice in water containing 6.7% acetic acid and 30% methanol, and then imaged in Odyssey CLx Infrared Imaging System (Li-COR). Post this, membranes were destained with 0.1 M NaOH containing 30% methanol. Membranes were blocked for 5 mins in EveryBlot Blocking Buffer (Bio-Rad Cat#12010020) and then incubated with the appropriate primary antibody overnight at 4^0^C. Membranes were washed three times the following day with 1X TBS (50mM Tris-HCl, 274mM NaCl, 9mM KCl) with 0.1% Tween 20 (TBST). Membranes were incubated with the corresponding secondary antibody for 1 hr at RT, washed thrice in TBST and then imaged in Odyssey CLx Infrared Imaging System. If membranes had to be stripped, they were done in 1X NewBlot™ IR Stripping Buffer (Licor Cat# 928-40028) for 20 mins at RT and re-probed again starting from the blocking step. Band intensities were quantified in Image Studio™ Software (Li-COR). The following primary antibodies were used in this study: Rubicon at 1:1000 (CST #D9F7; RRID: AB_10891617), MTMR2 at 1:2000 (sc-365184; RRID: AB_10708283), MTMR5 at 1:500 (sc-393488; RRID: AB_3097714), Parkin at 1:1000 (CST #2132; RRID: AB_10693040), VPS34 at 1:2000 (NB110-87320SS; RRID: AB_1199455), Mitofusin-2 at 1:1000 (sc-100560; RRID: AB_2235195), ATG7 at 1:1000 (ab133528; RRID: AB_2532126), ATG5 at 1:1000 (ab108327; RRID: AB_2650499), GAPDH at 1:1000 (ab9484; RRID: AB_307274), Actin at 1:1000 (Sigma-Aldrich MAB1501R; RRID: AB_2223041), Tubulin at 1:2000 (CST #2148; RRID: AB_2288042), LC3B at 1:1000 (ab48394; RRID: AB_881433), Ubiquitin at 1:1000 (sc-8017; RRID: AB_628423). The following secondary antibodies were used at 1:20,000 dilution: IRDye® 800CW Donkey anti-Mouse IgG (Li-cor #926-32212, RRID: AB_621847); IRDye® 680RD Donkey anti-Mouse IgG (Li-cor # 926-68072, RRID: AB_1095362); IRDye® 800CW Donkey anti-Rabbit IgG (Li-cor # 926-32213, RRID: AB_621848); IRDye® 680RD Donkey anti-Rabbit IgG (Li-cor # 926-68073, RRID: AB_10954442).

### Assays for autophagy, lysosomal function and mitophagy

**To assay for autophagy**, neurons were incubated with 50 nM DAP Red (Dojindo, D677-10) in fresh MM for 30 mins at 37^0^C. The culture medium was aspirated out and the neurons were washed with fresh MM. Neurons were then incubated with fresh MM containing 50 nM Lysotracker Green (ThermoFisher Scientific #L7526) for another 25-30 minutes at 37^0^C and imaged live in imaging media containing 50 nM Lysotracker Green.

**To test lysosomal function**, neurons were incubated with Magic Red Substrate (MR-RR2) (Biorad, ICT937) at a final dilution of 1:4000 in fresh MM for 15 mins at 37^0^C. Post incubation, neurons were imaged live in imaging media containing the Magic Red substrate at 1:4000 dilution.

**To assay for mitophagy**, neurons were incubated with 20 nM Mtphagy dye (Dojindo, MD01-10; labeled here as DMP Red) in fresh MM for 30 mins at 37^0^C. The culture medium was aspirated out and the neurons were washed twice with fresh MM. Neurons were then incubated with EtOH or 3 nM Ant A in MM for 1 hr and then incubated for additional 1.5 hrs in fresh MM without EtOH or Ant A. Lysotracker Green (ThermoFisher Scientific #L7526) at a final concentration of 50 nM was added, and the neurons were then incubated for a final 25-30 mins. The media was then aspirated out and imaged live in imaging media containing 50 nM Lysotracker Green.

### Ubiquitin enrichment assay

Neurons were treated with 10 µM MG132 for an additional 1 hr prior to adding 15 nM Ant A and incubating it for 2 hrs. Post incubation, neurons were lysed in lysis buffer containing 50 mM HEPES (pH=7.4), 1 mM EDTA, 1 mM MgCl_2_, 25 mM NaCl, 0.5% Triton X-100, 20μg/mL Leupeptin, DTT, 20 μg/mL TAME, 2 μg/mL Pepstatin A and 1 mM PMSF. Deubiquitination inhibitors provided with Ubiquitin Detection Kit (Cytoskeleton, BK161) were also added to the lysis buffer at 1X concentration. Post lysis of the neurons, lysates were spun at 17000x g for 10 mins, and the pellet was discarded. 5% of the supernatant was stored as input to be used for western blotting later. The immunoprecipitation assay was performed using Ubiquitin Detection Kit (Cytoskeleton, BK161) as mentioned in the manufacturer’s protocol with some modifications. Briefly, ubiquitination affinity beads and ubiquitination IP control beads were washed with 1XPBST twice and 1XPBS once, and then incubated with equal volumes of the lysate for 4 hrs at 4^0^C. Post incubation, beads were centrifuged at 5000x g for 4^0^C for 1 min. Beads were thrice washed with BlastR-2^TM^ wash buffer and centrifuged. After final wash and centrifugation, beads were incubated with Bead Elution Buffer for 5 mins at RT and then centrifuged in a spin column at 10000x g for 1 min to collect the immunoprecipitates. 2-mercaptoethanol was added to each sample which were then boiled at 95^0^C for 10 minutes to be used for western blotting.

### Live imaging and image analysis

When required, prior to imaging, neurons were labeled with 100 nM of Janelia Fluor 646 Halo Ligand (Promega, Cat: #GA112A) for 20 mins followed by a washout for another 20-30 mins. Neurons were imaged in imaging media containing HibernateE Brain Bits and supplemented with 33 mM D-glucose, 37.5 mM NaCl and 2% B27. Neurons were imaged live inside an environmental chamber at 37^0^C for 45-60 mins. Imaging was performed on a PerkinElmer UltraView Vox spinning disk confocal on a Nikon Eclipse Ti Microscope with an Apochromat 100x 1.49 N.A. oil-immersion objective and a Hamamatsu CMOS ORCA Fusion (C11440-20UP) camera with VisiView (Visitron).

To quantify colocalization between EGFP-Rubicon, LAMP1-Halo and mCherry-LC3 in the soma, punctae were identified manually from a single z-plane across channels and binarized. The number of colocalizing puncta was then calculated using the AND function in Fiji/ImageJ. For colocalization experiments with GFP-Rab7, LAMP-Halo and mCherry-Rubicon^WT^ or mCherry-Rubicon^CGHL^, cells areas from a single z-plane were thresholded and binarized using Yen thresholding in Fiji/ImageJ. Area of colocalization was then calculated using the AND function in Fiji/ImageJ.

To quantify DAP Red, Lysotracker numbers from a single z-plane the region of interest, i.e. the soma was identified. DAP Red and Lysotracker positive puncta were then segmented using Ilastik segmentation tool, a maching learning segmentation software. Images were binarized and thresholded, and the numbers calculated using the Analyze Particles option in Fiji/ImageJ. No. of colocalizing DAP Red and Lysotracker puncta were calculated using AND function.

To quantify no. of Magic Red positive punctae from a single z-plane, the region of interest, i.e. the soma was identified. Magic Red positive puncta were then segmented using Ilastik segmentation. Images were binarized and thresholded, and the numbers calculated using the Analyze Particles option in Fiji/ImageJ.

To measure mitophagolysosome nos. (i.e. colocalizing DMP Red and Lysotracker puncta), max projections of each channel from each image were generated. DMP Red and Lysotracker positive puncta from the soma were then segmented using Ilastik. Images were binarized and thresholded, and the no. of colocalizing DMP Red and Lysotracker puncta were calculated using AND function.

### Statistics

All statistical analysis were performed on Graphpad prism Version 10.1.1. All column graphs and superplots were generated on Graphpad prism Version 10.1.1. Data points from all biological replicates were tested for normal distribution using the Shapiro-Wilk normality test. Each data point on a column graph represents one biological replicate. For superplots, each biological replicate has been represented by colored shapes with black outlines, and its corresponding technical replicates represented with the same translucent color and shape. All analysis was performed on at least three biological replicates per condition. Data are represented as mean and the standard error of mean (S.E.M.).

## DATA AVAILABILITY

Source data will be made publicly available on Zenodo upon formal acceptance of the manuscript.

## ACKNOWLEDGEMENTS

We thank Julia F. Riley for sharing astrocyte samples, and Mariko Tokito and Karen Jahn for technical assistance. We also thank Dr. Sierra Palumbos, Elizabeth R. Gallagher, Dr. Juliet Goldsmith, Dr. Kaya Matson, and all other members of the Holzbaur laboratory for insightful discussions and valuable feedback. We thank Dan A. Tudorica and Prof. James H. Hurley (UC, Berkeley) for gifting us the EGFP-Rubicon clone. We also thank Dr. Dorotea Fracchiolla for help with project management.

This work was supported by the Michael J. Fox Foundation (MJFF) (MJFF-021130, MJFF-15100, and MJFF-019411 to E.L.F. Holzbaur. The E.L.F. Holzbaur laboratory is funded by the joint efforts of The MJFF and the Aligning Science Across Parkinson’s initiative. MJFF administers the grant ASAP-000350 on behalf of ASAP and itself.

