## Supplemental Figures for "Mitochondrial damage triggers concerted degradation of negative regulators of neuronal autophagy"

**Supplemental S1**

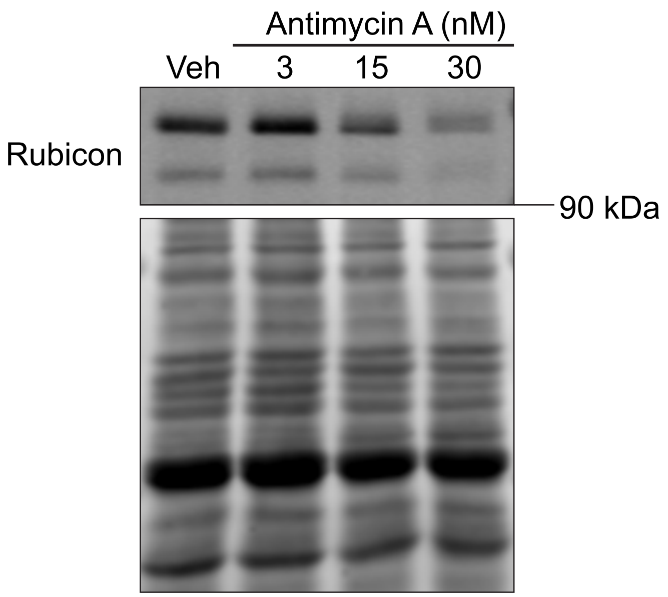

**Supplemental Figure S1:** Mitochondrial damage induces parallel degradation of both the putative isoforms of Rubicon. Representative western blot from lysates of WT cortical neurons treated with vehicle (EtOH) or with increasing concentrations of Ant A (3 nM, 15 nM or 30 nM) for 2 hrs and probed for Rubicon. A doublet is seen on the immunoblot.

**Supplemental S2**

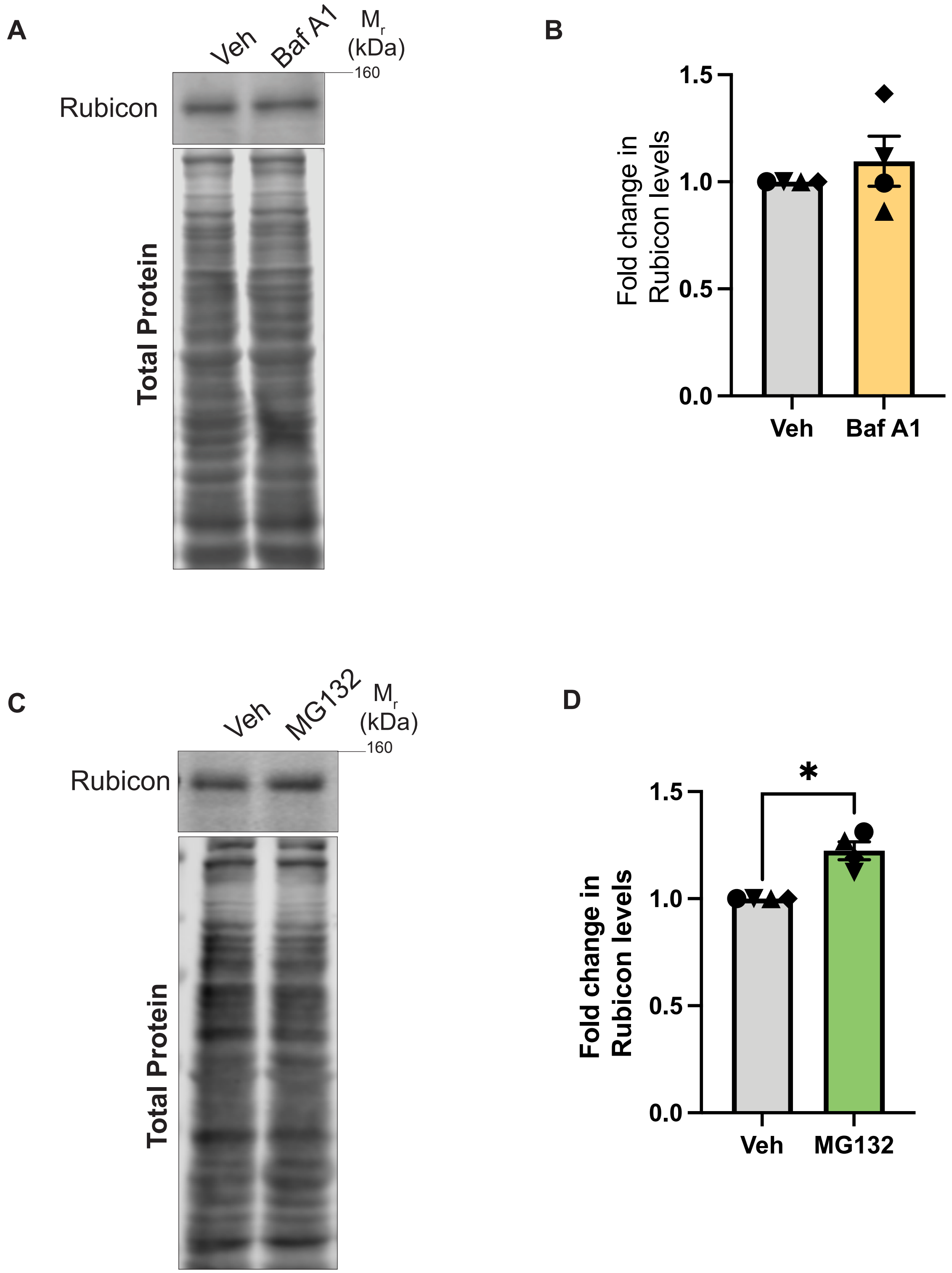

**Supplemental Figure S2: (A, B)** Rubicon levels are unaltered upon blocking basal autophagy in HeLa cells. (A) Representative western blot from lysates of HeLa cells treated with vehicle (DMSO) or 500 nM Baf A1 for 3 hrs. (B) Rubicon band intensity normalized to total protein intensity from lysates of HeLa cells treated with DMSO or Baf A1 for 3 hrs. **(C, D)** Rubicon levels are not regulated by the proteasome in HeLa cells under basal conditions. (C) Representative western blot from lysates of HeLa cells treated with vehicle (EtOH) or 10 µM MG132 for 3 hrs. (D) Rubicon band intensity normalized to total protein intensity from lysates of HeLa cells treated with EtOH or MG132 for 3 hrs. All panels: N=4 experiments, Mann-Whitney test, error bars indicate S.E.M.

**Supplemental S3**

A

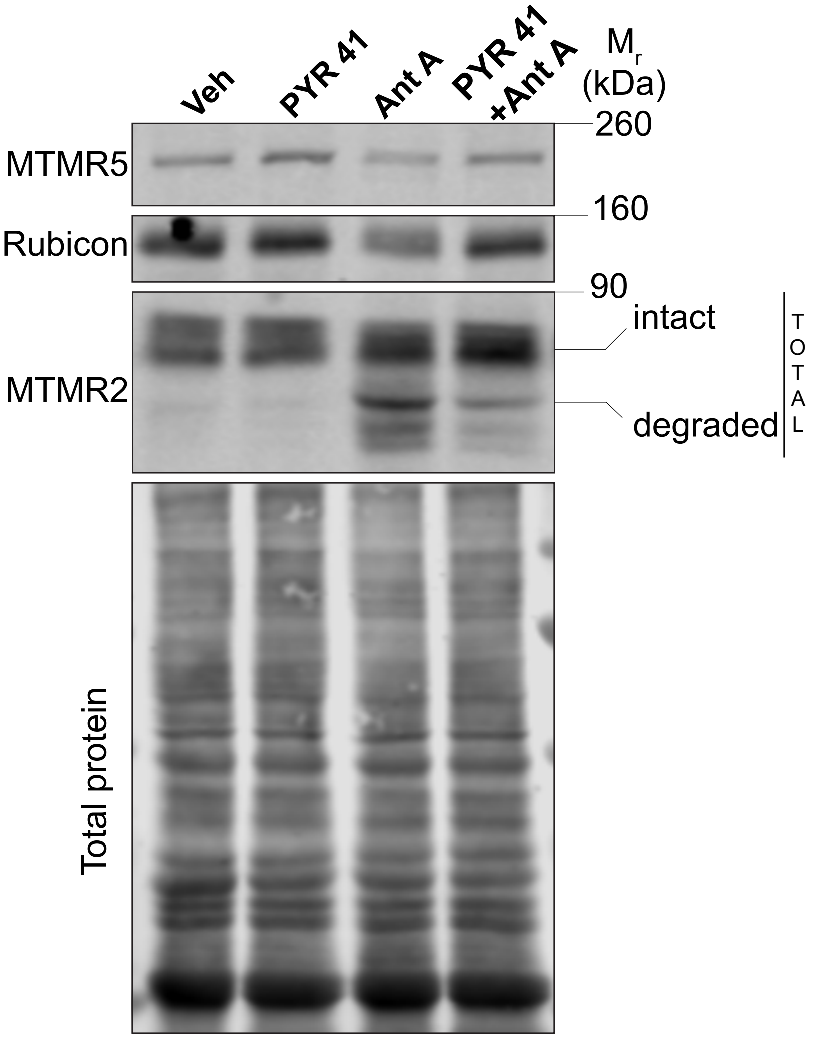


B

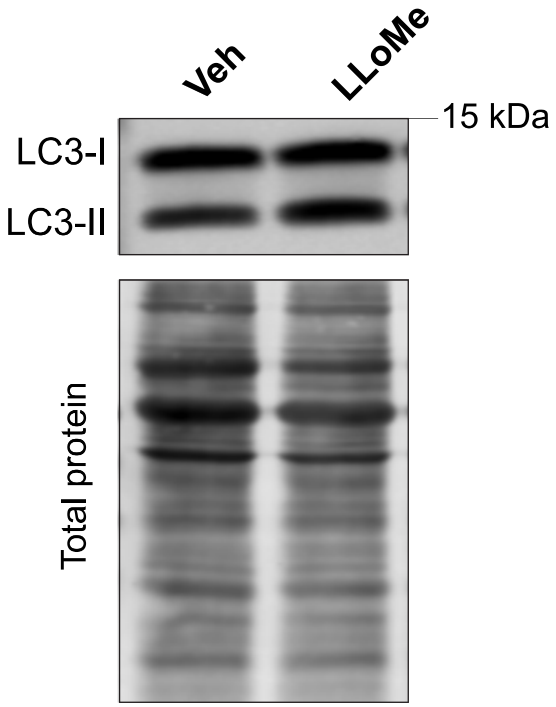

**Supplemental Figure S3:** (A) Blocking Ub-E1 activity by PYR 41 suppresses the degradation of MTMR5/2 and Rubicon during mitochondrial damage. Representative western blot from lysates of WT embryonic cortical neurons treated with vehicle (DMSO) or 10 μM PYR 41 for 1 hr followed by an additional treatment with vehicle (EtOH) or 15 nM Ant A for 2 hrs. Experiments were performed thrice and one of the replicates is shown here. **(**B) Lysosomal damage induced by LLoMe results in accumulation of LC3-II in neurons. Representative western blot from lysates of WT embryonic cortical neurons treated with vehicle (EtOH) or 1 mM LLoMe for 2 hrs and probed for LC3. Experiments were performed twice and one of the replicates is shown here.

### Supplemental S4

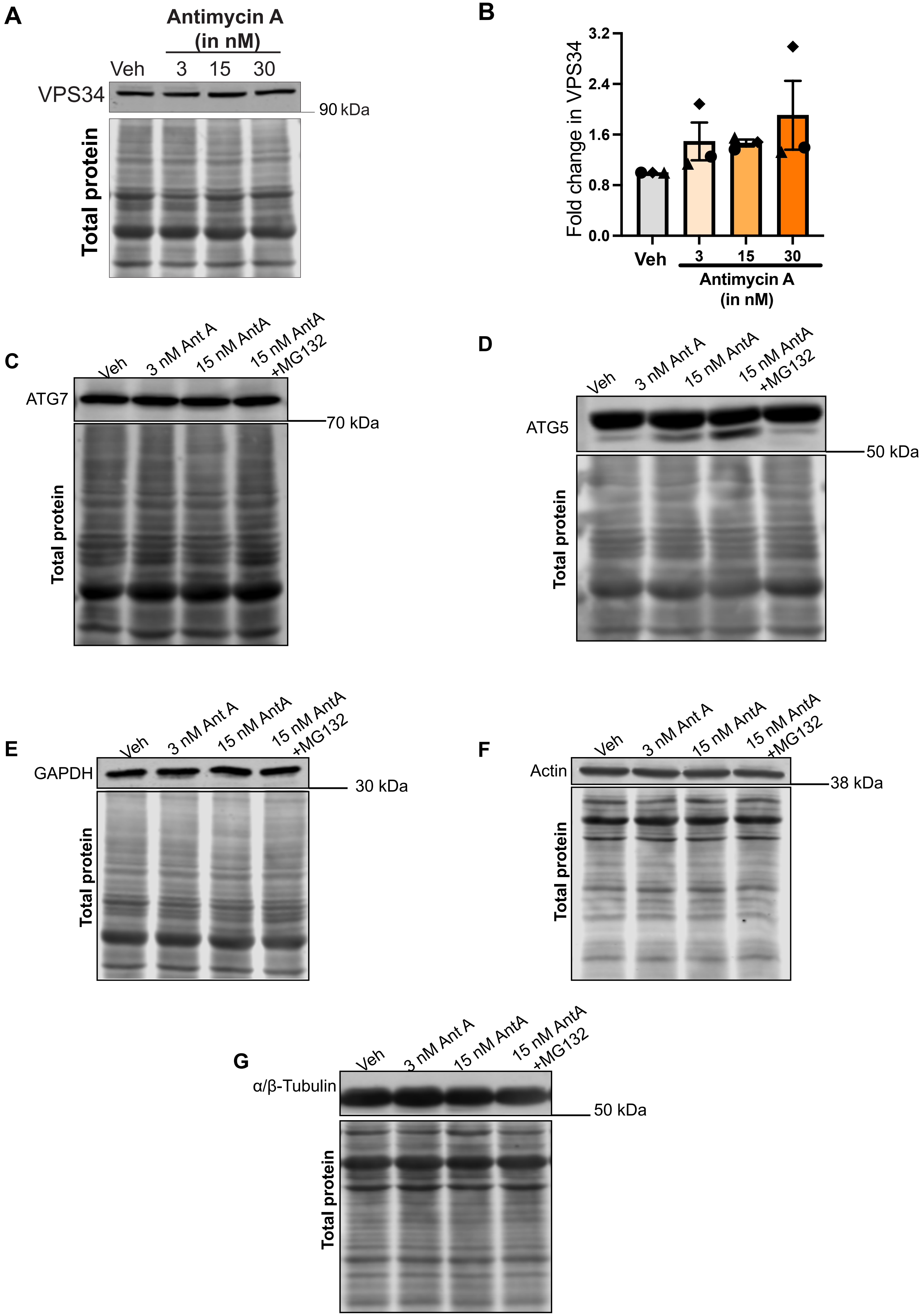

**Supplemental Figure S4: (A-B)** Levels of VPS34, a positive regulator of autophagy, are unchanged during mitochondrial damage. (A) Representative western blot from lysates of WT cortical neurons treated with vehicle (EtOH), or with increasing concentrations of Ant A (3 nM, 15 nM or 30 nM) for 2 hrs (vehicle=EtOH) and probed for VPS34. (B) VPS34 band intensity normalized to total protein from lysates of WT neurons treated with EtOH or increasing concentrations of Ant A (One way ANOVA with Dunnett's multiple comparison test, ns=not significant, error bars indicate S.E.M.). **(C-G)** Positive regulators of autophagy (ATG7, ATG5) and house-keeping proteins (GAPDH, Actin, α/β-tubulin) are not subjected to proteasome mediated degradation upon mitochondrial stress. Representative western blots from lysates of WT cortical neurons treated with vehicle (EtOH), 3 nM Ant A, 15 nM Ant A or 15 nM Ant A+ 10 μM MG132. MG132 was added 1 hr prior to an additional treatment with vehicle (EtOH) or 15 nM Ant A for 2 hrs. Blots were probed for: (C) ATG7, (D) ATG5, (E) GAPDH**, (**F) Actin**, (**G) α/β-tubulin**.** All panels: Experiments were performed thrice and one of the replicates is shown here.

### Supplemental S5

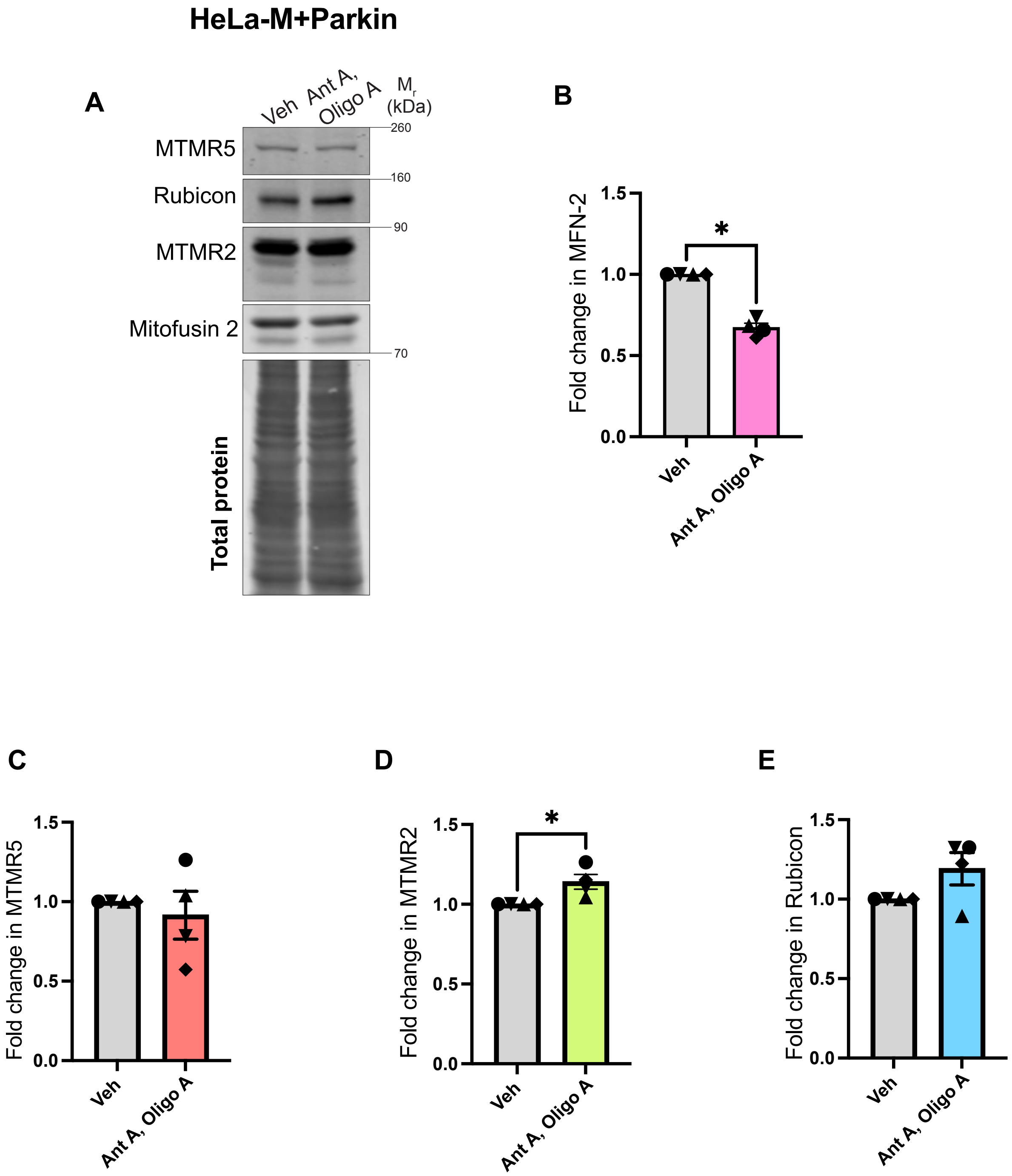

**Supplemental Figure S5: (A-E)** HeLa cells activate Pink1/Parkin-dependent mitophagy in response to mitochondrial damage but not MitoSR. (A) Representative western blot from lysates of HeLa cells transfected with an untagged construct of human Parkin and then treated with 10 µM Ant A, 10 µM Oligo A or vehicle (EtOH, DMSO) for 5 hrs. (B) Fold change in Mitofusin-2 levels upon treatment of Parkin-transfected HeLa cells with 10 µM Ant A, 10 µM Oligo A as compared to with vehicle. **(**C) Fold change in MTMR5 levels upon treatment of Parkin-transfected HeLa cells with 10 µM Ant A, 10 µM Oligo A as compared to with vehicle. **(**D) Fold change in MTMR2 levels upon treatment of Parkin-transfected HeLa cells with 10 µM Ant A, 10 µM Oligo A as compared to with vehicle. **(**E) Fold change in Rubicon levels upon treatment of Parkin-transfected HeLa cells with 10 µM Ant A, 10 µM Oligo A as compared to with vehicle. All panels: N=4 experiments, *$p<0.05$, Mann-Whitney test, error bars indicate S.E.M.

### Supplemental S6

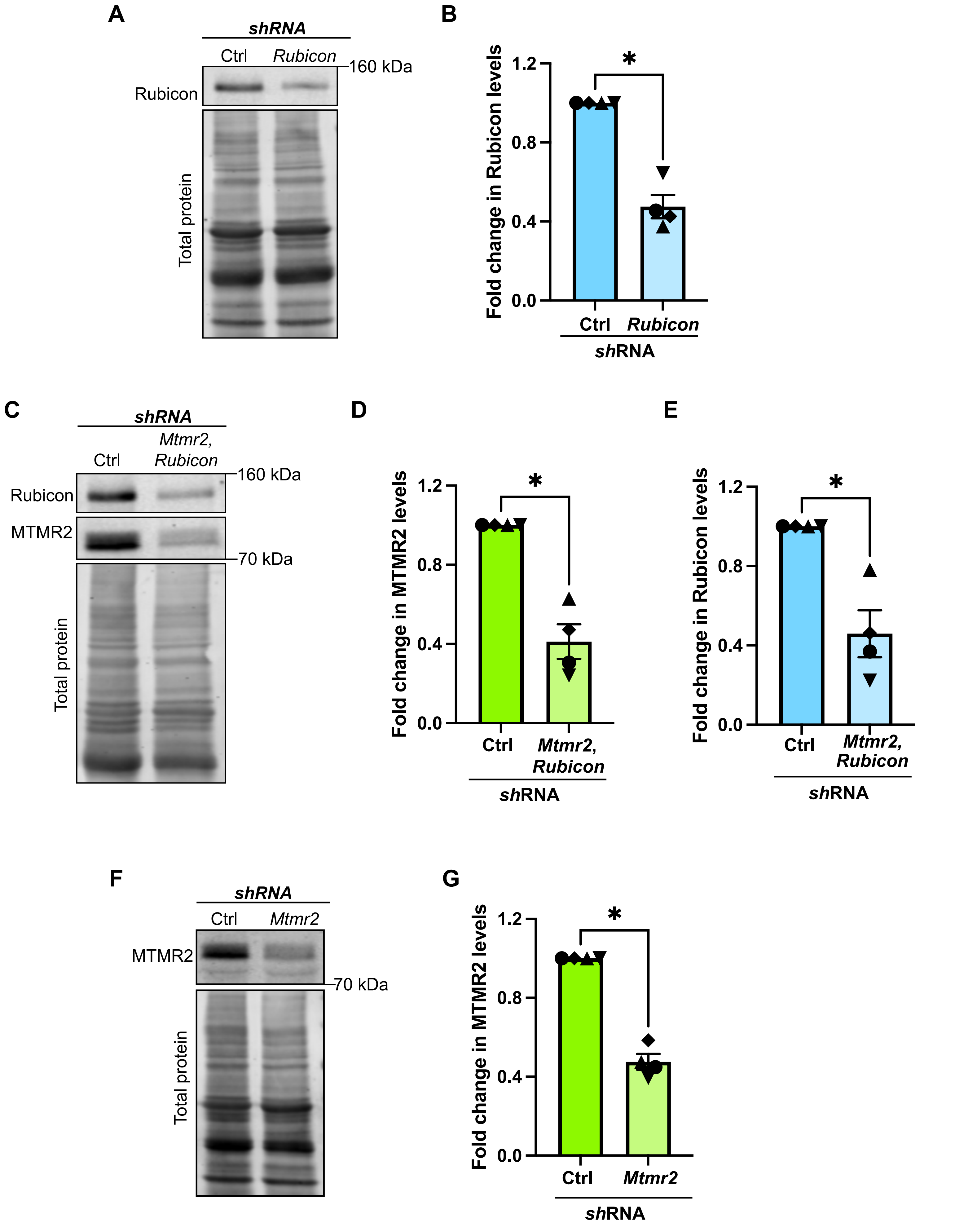

**Supplemental Figure S6: (A-B)** Knockdown of Rubicon in WT cortical neurons to determine its role in regulating basal autophagy and mitophagy. **(**A) Representative western blot from lysates of WT cortical neurons nucleofected with control or *Rubicon* shRNA plasmid, and probed for Rubicon. (B) Rubicon band intensity normalized to total protein upon knockdown of Rubicon in WT cortical neurons. **(C-E)** Knockdown of MTMR2 and Rubicon in WT cortical neurons to determine their role in regulating mitophagy. **(**C) Representative western blot from lysates of WT cortical neurons nucleofected with control or *Mtmr2* and *Rubicon* shRNA plasmids, and probed for MTMR2 and Rubicon. (D) MTMR2 band intensity normalized to total protein upon knockdown of Rubicon and MTMR2 in WT cortical neurons. (E) Rubicon band intensity normalized to total protein upon knockdown of Rubicon and MTMR2 in WT cortical neurons. **(F-G)** Knockdown of MTMR2 in WT cortical neurons to determine its role in regulating mitophagy. **(**F) Representative western blot from lysates of WT cortical neurons nucleofected with control or *Mtmr2* shRNA plasmid, and probed for MTMR2. (G) MTMR2 band intensity normalized to total protein upon knockdown of MTMR2 in WT cortical neurons. All panels: N=4 experiments, *$p<0.05$, Mann-Whitney test, error bars indicate S.E.M.
